# Replaying the evolutionary tape to investigate subgenome dominance in allopolyploid *Brassica napus*

**DOI:** 10.1101/814491

**Authors:** Kevin A. Bird, Chad Niederhuth, Shujun Ou, Malia Gehan, J. Chris Pires, Zhiyong Xiong, Robert VanBuren, Patrick P. Edger

**Author notes:** Correspondence to P.P.E., R.V., or Z.X.

## Abstract

Interspecific hybridization and allopolyploidization merges evolutionarily distinct parental genomes (subgenomes) into a single nucleus. A frequent observation is that one subgenome is “dominant” over the other subgenome, having a greater number of reatined duplicate genes and being more highly expressed. Which subgenome becomes dominantly expressed in allopolyploids remains poorly understood. Here we “replayed the evolutionary tape” with six isogenic resynthesized *Brassica napus* (rapeseed) allopolyploid lines and investigated subgenome dominance patterns over the first ten generations. We found that the same subgenome was consistently more dominantly expressed in all lines and generations. Furthermore, DNA methylation differences between subgenomes mirrored the observed gene expression bias towards the *Brassica oleracea* derived ‘C’ subgenome in all lines and generations. These differences in gene expression and methylation were also found when comparing the progenitor genomes, suggesting subgenome dominance is related to inherited parental genome differences rather than a byproduct of allopolyploidization. Gene network analyses indicated an enrichment for network interactions and several biological functions for ‘C’ subgenome biased pairs, but no enrichment was observed for ‘A’ subgenome biased pairs. These findings demonstrate that “replaying the evolutionary tape” in allopolyploids results in repeatable and predictable subgenome expression dominance patterns based on preexisting genetic differences among the parental species. These findings have major implications regarding the genotypic and phenotypic diversity observed following plant hybridization in both ecological and agricultural contexts.

## Introduction

Hybridization among closely related species is a widespread and recurrent evolutionary process (Arnold and Meyer 2006; Mallet 2007; Soltis and Soltis 2009). By merging the genomes of independently evolved species into a single nucleus, hybridization creates a unique opportunity for immense variability that natural selection can act upon in subsequent generations (Anderson and Stebbins 1954; Rieseberg et al. 2003). Furthermore, hybridization is known to produce transgressive phenotypes, including heterosis, and novel phenotypic variation not observed in the parents (Pires et al. 2004; Dittrich-Reed and Fitzpatrick 2013). However, the hybridization of highly diverged genomes, particularly those with different base chromosome numbers, can also lead to chromosome pairing issues during meiosis which greatly reduces fertility. Proper bivalent pairing of homologous chromosomes in such interspecific hybrids can be restored through whole genome duplication (i.e. polyploidization) resulting in the formation of an allopolyploid species (Charron et al. 2019). This may in part explain the high prevalence of polyploidy across flowering plants (Leitch and Leitch 2008; Van de Peer et al. 2009).

Genome-scale analyses of recent and ancient allopolyploids led to the discovery that one of the parental species’ genomes (also known as subgenomes) often exhibits greater gene retention (Thomas 2006), more tandem gene duplications (Edger et al. 2019), higher gene expression (Schnable et al. 2011) and lower DNA methylation (Woodhouse et al. 2014). Collectively this phenomenon is referred to as ‘subgenome dominance’. A previous study demonstrated that subgenome dominance at the gene expression level occurs at the moment of interspecific hybridization and increased over subsequent generations in the allopolyploid (Edger et al. 2017). This finding agrees with theoretical work of transcription factor binding and regulatory mismatch that predicts increasing subgenome dominance over generations in newly established allopolyploids (Bottani et al. 2018). Preexisting differences between parental genomes has been shown to influence these observed subgenome dynamics in an allopolyploid (Buggs et al. 2014; Kryvokhyzha et al. 2019). For example, analyses of diverse allopolyploids have revealed that gene expression differences among subgenomes mirrors differences in transposable element (TE) densities in flanking regions surrounding genes (Freeling et al. 2012; Cheng et al. 2016; Edger et al. 2019). These findings collectively suggest that subgenome dominance may be largely predetermined based on subgenome differences in certain genomic features including TE densities.

Given that gene expression level dominance occurs instantly following the initial hybridization event (Edger et al. 2017), resynthesized allopolyploids are the ideal system to investigate the establishment and escalation of subgenome dominance. Few studies have used multiple independently derived resynthesized allopolyploids to investigate subgenome dominance (Chagué et al. 2010; Combes et al. 2015; Hao et al. 2017; Wu et al. 2018; Gaebelein et al. 2019; Li et al. 2019). It remains unclear the extent to which the emergence of subgenome expression dominance is a result of pre-existing characteristics of the diploid progenitors or due to independent and non-recurrent events during polyploid formation. In other words, will multiple independently established allopolyploids consistently exhibit the same patterns of subgenome expression dominance (e.g. towards the same subgenome)?

Here we analyzed subgenome dominance in six independent resynthesized allopolyploid *Brassica napus* (2n=4x=38) lines formed by hybridizing two doubled haploid parents from the progenitor species *Brassica rapa* (AA; 2n=2x=20) and *Brassica oleracea* (CC; 2n=2x=18) (Song et al. 1995). The crop *B. napus* was formed between 7,500 and 12,500 years ago and is widely grown present-day as an oilseed crop (rapeseed), vegetable fodder crop (rutabaga) and vegetable crop (siberian kale) (Chalhoub et al. 2014; An et al. 2019). The strengths of the *B. napus* polyploid system include not only having high-quality reference genomes for both diploid progenitors and *B. napus*, but also being closely related to the model plant *Arabidopsis thaliana,* allowing for the integration of diverse genomic and bioinformatic resources (Cheng et al. 2012; Chalhoub et al. 2014; Parkin et al. 2014; Koenig and Weigel 2015). Furthermore, a previous analysis of the *B. napus* reference genome identified a greater number of retained genes in the *B. oleracea* (C) subgenome compared to *B. rapa* (A) subgenome (Chalhoub et al. 2014). This is consistent with patterns observed in older allopolyploids that exhibit subgenome dominance - dominant subgenome retaining a greater number of genes (Bird et al. 2018). Lastly, because the resynthesized *B. napus* lines were made with doubled haploids, each of the independent lines started out genetically identical (Ren et al. 2017). This permitted us to examine and compare the establishment of subgenome dominance across independently derived polyploid lines without the added influence of allelic variation segregating between different lines. We examined each of the six resynthesized polyploid lines with RNA-seq and Bisulfite-seq data to characterize gene expression and methylation differences between high confidence homoeologs over the first ten generations. This permitted us to assess the variability of subgenome dominance during the earliest stages following allopolyploid formation.

## Results

### Homoeolog Expression Bias

This population of resynthesized polyploids provided a unique opportunity to examine if the same subgenome would repeatedly exhibit subgenome expression dominance. Gene expression in leaves was surveyed using RNAseq in sixteen of the eighteen resequenced individuals (six lines and three generations). Library construction failed for two individuals, thus these were not able to be included in this analysis. However, all six lines were able to be investigated in these sets of analyses. Samples were aligned to an *in silico* polyploid reference genome. We restricted gene expression analyses to genomic regions with balanced gene dosage (2:2; AA:CC) identified using genome resequencing data for each individual to reduce the confounding factor of dosage changes in regions that have undergone homoeologous exchange. Expression patterns of the six lines were also compared to the parental *B. rapa* and *B. oleracea* genotypes to test if expression differences may exist among the diploid progenitors.

The mean expression bias (Log2 FoldChange BnC expression /BnA expression) for homoeologs in balanced (2:2) regions ranged from 0.12 to 1.16 (median −0.50 to 0.96), with 15 of 16 individuals having mean expression bias significantly greater than 0 (One-way Wilcoxian-Mann-Whitney test, p<2.2e^-16; Fig 1, FigS1-5). These results suggest a transcriptome-wide bias in favor of the C subgenome, however the magnitude of the expression bias was smaller than observed previously in other allopolyploids (Edger et al. 2017). The homoeolog expression bias between the parents was also significantly greater than 0 in these balanced regions. Comparing bias difference between the parental lines and the synthetic polyploids revealed that only 5/16 were significantly different from the parents (two way Wilcoxian-Mann-Whitney test, p < 0.001).

**Figure 1:**
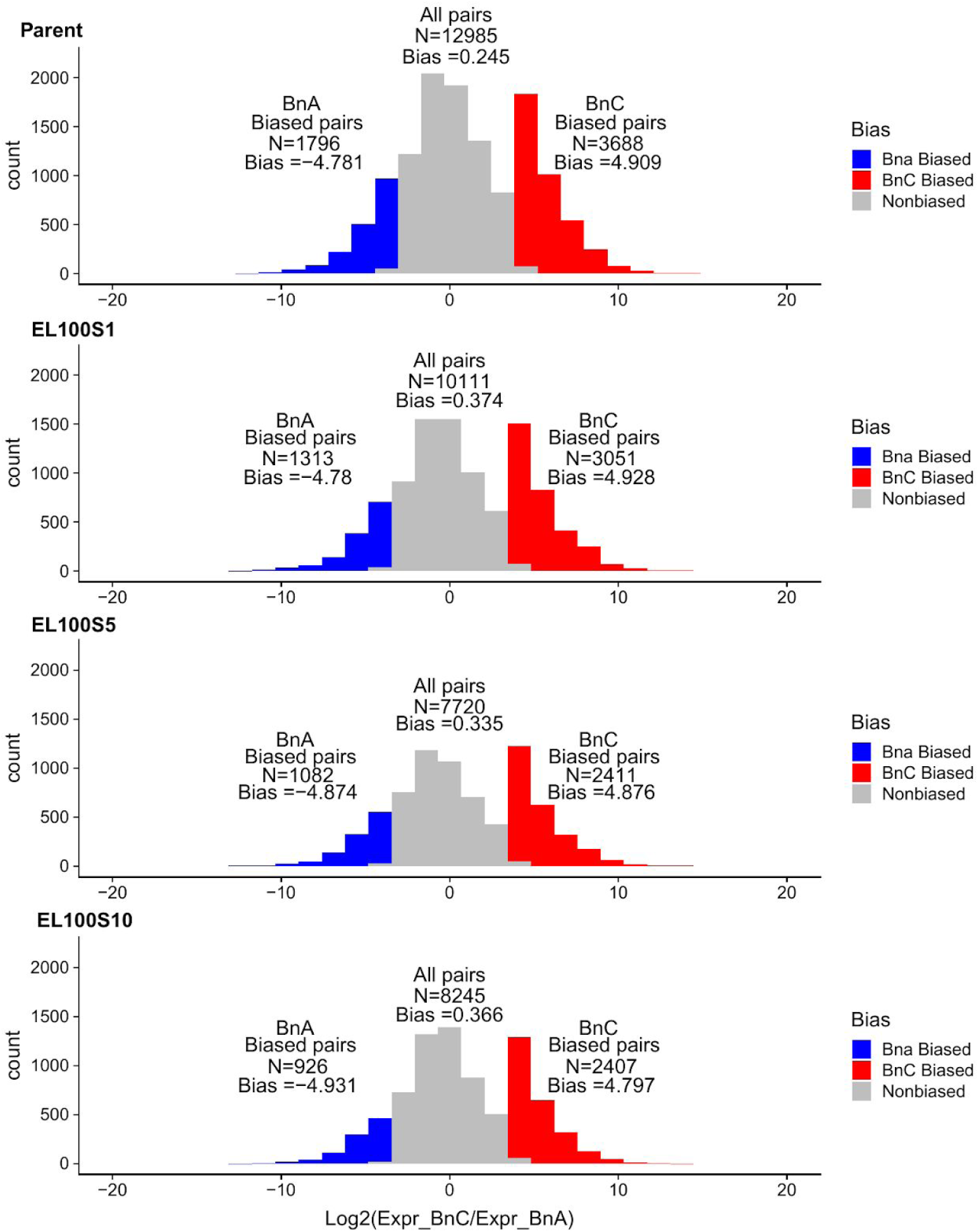
**Homoeolog Expression Bias**. Distribution of homoeolog bias in the parent and three generations of line EL100, red regions indicate BnC biased homoeologs with log2 expression fold change greater than 3.5 and blue regions indicate BnA biased homoeologs with log2 expression fold change less than −3.5.

We examined homoeolog pairs that were biased toward either subgenome (Log2 fold change > |3.5|) and found there were significantly more BnC biased homoeolog pairs than expected in all individuals (χ^2^ test, χ^2^ > 170, df=1, p < 2.2e^16; Figure 1; Figure S1-5; Table S1). This indicates that there is a bias in homoeolog expression on a pair-by-pair basis towards the BnC subgenome. This BnC homoeolog bias was consistent across generations. We also found that homoeolog bias in the synthetic polyploids was significantly different than existing expression bias in the parents for 7 of 16 individuals (χ^2^ test, χ^2^ =>12.459, df =2, p < 0.003125; Figure 1; Figure S1-5; Table S2). In 5 of those 7 individuals there were more BnC biased homoeologs than expected based on expression biases of the progenitor genomes. A bimodal distribution was observed when comparing A and C subgenome expression; the rightmost distribution being largely due to homoeolog pairs with a lack of BnA expression.

Next we investigated whether individual gene pairs were biased in the same direction across the six lines. Due to the stochastic nature of HEs, dosage of a gene pair may differ between lines. To adjust for this, we first looked only at genes found in 2:2 dosage for all lines in a generation, resulting in 6917, 3574, and 2252 homoeologous pairs, for generations S1, S5, and S10 respectively. A majority of C biased Gene pairs were biased towards the C subgenome in all synthetic lines for each generation and the parents (S1=1806 [75%], S5=772 [71%], and S10=602 [70%]; Fig 2a-c). In the first generation, roughly 36 gene pairs (1.5%) were uniquely dominant in only the parents and 32 (1.3%) were dominant in all six synthetic lines but not the parent (Fig 2a). In the fifth and tenth generation, there was a similar number of BnC dominant homoeologs that were dominant in only the parents (S5=17 [1.5%], S10=14 [1.6%]) and dominant in all six lines but not the parent (S5=17 [1.5%], S10=13 [1.5%] (Fig 2c). Similar patterns were observed for BnA biased homoeologs with most genes showing similar bias in all lines across generations and the parents (S1 = 698 [61%], S5 = 401 [58%], S10 = 221 [51%]; respectively; Fig S6a-c), and a consistently low number of genes biased in all six lines but not the parents in each generation (S1 = 28 [2.5%], S5 = 15 [2.2%], S10 = 10 [2.3%]) and only those biased in the parents (S1 = 41 [3.6%], S5 = 22 [3.2%], S10 = 18 [4.1%]; Fig S6a-c).

**Figure 2.**
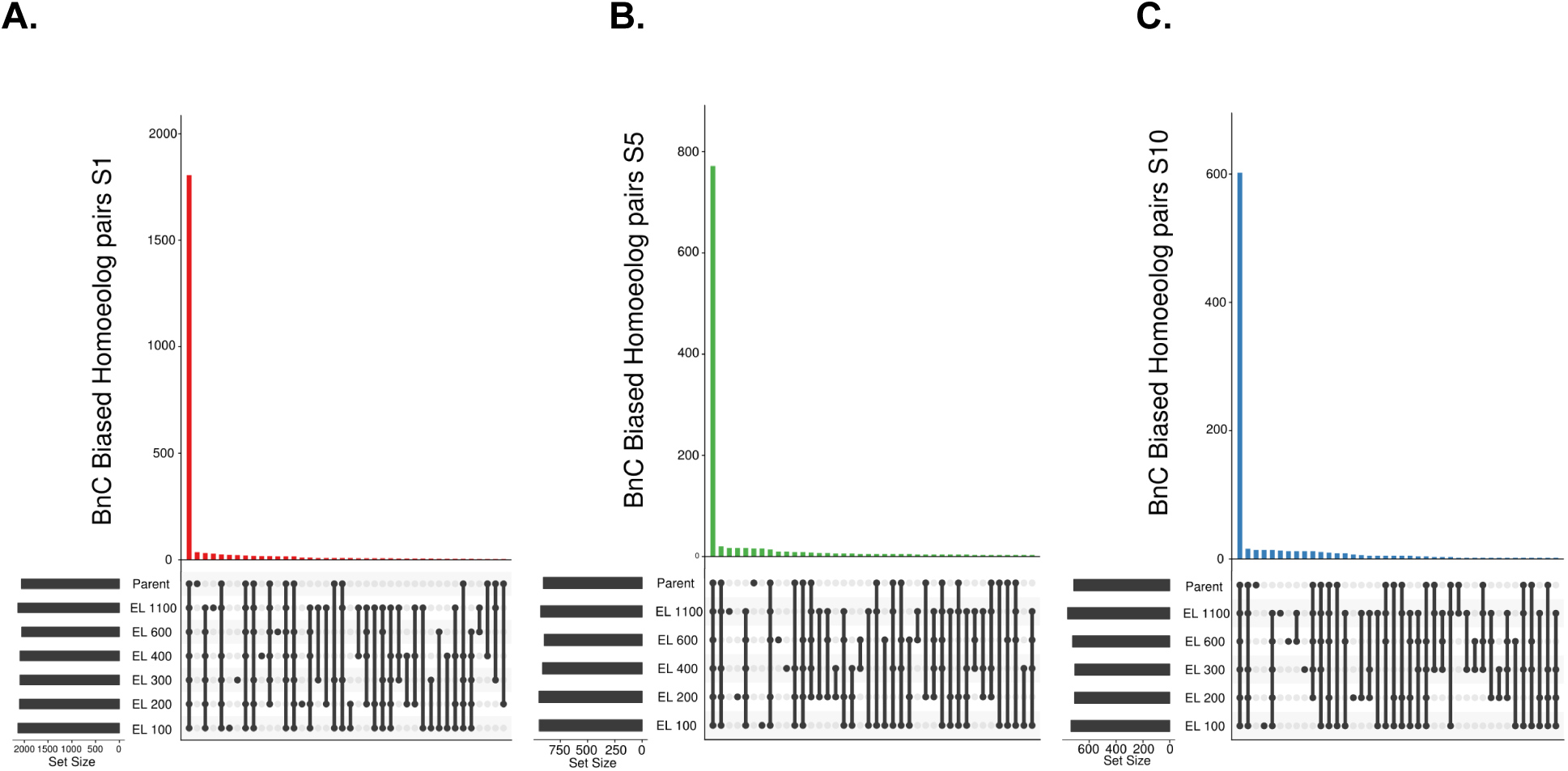
**Common shared biased homoeolog pairs**: Upset plot of how biased homoeologs pairs for BnC biased for generation 1 **(a)**, 5 **(b),** and 10 (**c)** are shared among all six lines for the three sampled generations. These analysis was restricted only to homoeolog pairs in 2:2 balance in all 6 lines

Lastly, we performed gene ontology (Ashburner et al. 2000), pathway (Kanehisa and Goto 2000) and network (Szklarczyk et al. 2017) enrichment analyses to determine if BnC and BnA biased gene pairs were enriched with certain biological functions. Placing biased homoeologs in a network context allowed us to determine the extent to which the genes interact with each other, the average number of connections a gene has with other genes (average node degree) and the extent to which genes are clustered together in the network. In all generations, BnC biased gene pairs showed enrichment for interactions in the PPI network in generations S1, S5, and S10 (p<e^-16) with high average node degree (S1=20.6, S5=12.1, S10=7.99) and clustering coefficient (S1=0.386, S5=0.385, S10=0.385). GO terms for BnC biased gene pairs were highly enriched with core metabolic, including photosynthesis, organellar, and ribosomal functions. BnC biased gene pairs were also significantly overrepresented in KEGG pathways annotated for amino acid biosynthesis, MAPK signalling and photosynthesis (Table S3-5). The enrichment for organellar functions for BnC biased pairs may be expected given that the C subgenome is the maternal progenitor of the resynthesized lines. This suggests that subgenome dominance may, in part, be related to maintenance of balanced nuclear-organellar interactions. On the contrary, BnA biased gene pairs showed no enrichment for any GO terms or KEGG pathways. However, there is still observed enrichment for interactions in the PPI network for BnA biased gene pairs in generations S1 (p=9.75e^-07) and S5 (p=0.00681), but not generation S10 (p=0.16). Lower network statistics were also observed for average node degree (S1=1.13, S5=0.912, S10=0.338) and average clustering coefficient (S1=0.270, S5=0.235, S10=0.209) for BnA biased gene pairs. In summary, subgenome expression dominance in *Brassica napus* are biased towards a set of highly interconnected BnC genes that are enriched for a wide variety of biological processes.

### DNA methylation

Cytosine methylation is involved in defining regions of chromatin, silencing transposons, maintaining genome integrity, and can affect gene expression. DNA methylation itself is shaped by factors such as gene expression and the underlying sequence (Niederhuth and Schmitz 2017). To understand how DNA methylation evolves following polyploidy and may contribute to subgenome dominance, DNA methylation was assessed using whole-genome bisulfite sequencing (WGBS) in the parents and allopolyploids in this population (Cokus et al. 2008; Lister et al. 2008). In plants, three sequence contexts are generally recognized for DNA methylation, depending on second and third bases downstream of the methylated cytosine: CG (or CpG), CHG, and CHH (where H= A, T, or C). Methylation of these contexts are established and maintained by different molecular pathways and their associations with gene expression can differ based on the pattern of DNA methylation, hence they are typically analyzed separately (42).

In all methylation contexts the C-subgenome progenitor showed higher methylation levels than the A-subgenome progenitor.

When the entire genome is analyzed as a whole, CG methylation in the resynthesized lines typically fell between the two parents and showed little change in genes and Long-Terminal Repeat (LTR) retrotransposons, regardless of generation (Fig S7a, S8a). More variance in total methylation was observed for the non-CG contexts. In the first generation, CHG methylation was comparable to either parent. In generations five and ten, CHG methylation became increasingly variable (Fig S7b, Fig S8b). CHH methylation levels were even more striking. A slight increase in CHH methylation was observed in generation one and this trend became more pronounced in generations five and ten, with methylation levels surpassing the highest parent (Fig S7c, S8c).

Analyzed by subgenomes, methylation in the resynthesized lines was lower on the A subgenome than the C subgenome (Fig S9-S12). Additionally, a number of individuals showed increased CG methylation in genes and flanking regions for BnC genes, but little difference is observed for BnA genes (Fig S9a,c). In genic regions on the BnC subgenome, CG methylation increase occurred primarily for non-syntenic genes (Fig S11c), however in flanking regions of genes, CG methylation increases occurred for both syntenic and non-syntenic genes (Fig S11c S12c). For CHG sites, methylation levels in the first generation samples again showed lower levels both in the bodies and in flanking regions of genes and LTR retrotransposons, and increased in subsequent generations of some lines in both the BnA and BnC subgenome (Fig S10b,e). CHH methylation again showed the most striking differences. First generation allopolyploids showed little difference from the parents, when all genes were examined (Fig S8c). However, when examined by subgenome, BnC genes show increased CHH methylation in flanking regions (Fig S9c), while BnA genes did not (Fig S9f). In subsequent generations, CHH methylation increased in both LTR retrotransposons and gene flanking regions (Fig S9c,f; S10c,f). For BnC genes, CHH methylation of gene flanking regions and LTR retrotransposons increased beyond the initial increase in the first generation (Fig S9f, S10f). Increased CHH methylation was not found in flanking regions of BnA genes (Fig S9c) but was observed to a lesser extent in LTR retrotransposons on the BnA subgenome (Fig S10c). These changes in flanking CHH methylation occurred for both syntenic and non-syntenic genes (Fig S11c,f; S12c,f). Additionally, we saw higher methylation of LTR retrotransposons in the dominant subgenome (BnC) than the submissive subgenome (BnA) (Fig S10c,f).

To investigate the potential roles and patterns of methylation related to subgenome dominance, we again focused on homoeologs identified in 2:2 dosage with biased expression in the previous analyses. For all lines there was higher methylation in all contexts in homoeologs on the BnC subgenome than the BnA subgenome. Additionally all lines showed higher methylation levels 2kb up- and downstream of the gene body in BnC biased homoeolog pairs in the CG, CHG, and CHH methylation context (Fig 3). Furthermore, there was a notable difference between BnA and BnC biased homoeologs at the transcription start site (TSS’), with BnC biased homoeologs showing lower CG methylation levels (Fig 3). Similarly BnC biased homoeologs showed lower CHH and CHG gene body methylation levels.

**Figure 3:**
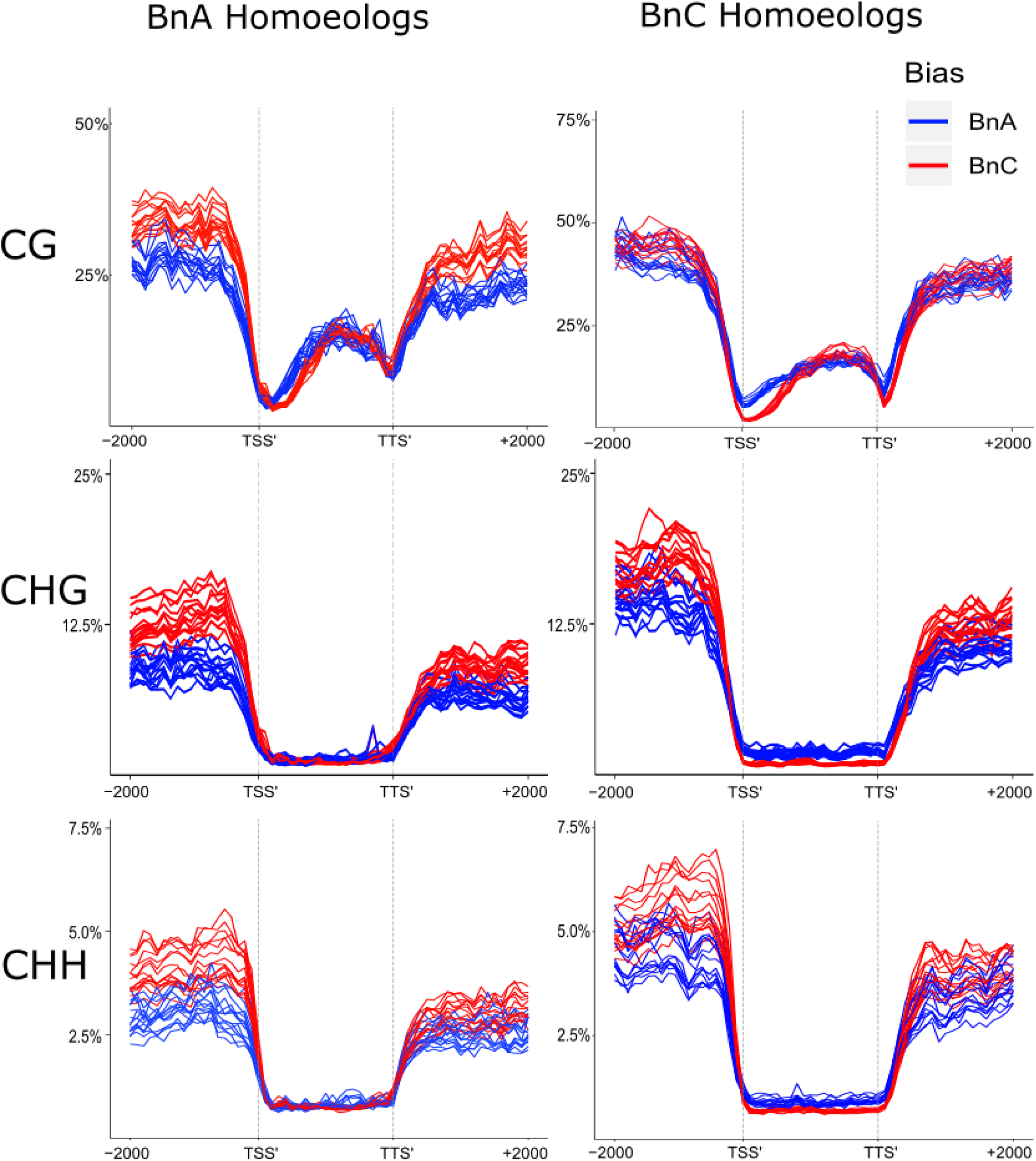
**DNA Methylation Bias.** Average weighted methylation for homoeolog pairs in 2:2 dosage for all 16 lines assessed using Bisulfite-seq. BnC biased expressed homoeologs in red and BnA biased expressed homoeologs in blue. Results are shown for methylation in CG, CHG, and CHH contexts. Methylation levels are shown for the transcription start site (TSS), gene body, transcription termination site (TTS) and 2kb up- and downstream of the TSS and TTS.

## Discussion

Our findings suggest that “replaying the evolutionary tape” yields nearly identical subgenome expression dominance patterns in these six *Brassica napus* resynthesized lines. By leveraging independent resynthesized polyploid lines from the same doubled-haploid parents, we were able to assess subgenomic expression and methylation patterns without confounding bias from standing allelic variation. We discovered significant expression bias favoring the BnC subgenome and a significantly higher number of BnC biased homoeologous pairs in all six lines that was consistently maintained over the ten generations. Furthermore, we discovered that these expression differences between homoeologs in the polyploids mirrored pre-existing expression differences among the diploid parents. We also identified methylation differences between homoeolog pairs that reflected observed gene expression differences. Similar to gene expression differences, these methylation differences were largely pre-existing in the parental diploid epigenomes. Finally we identified transgressive hypermethylation at CHH sites on genes and LTR transposable elements across both subgenomes, likely as a response to “genomic shock” following interspecific hybridization. These results suggest that subgenome expression dominance in an allopolyploid is predictable and repeatable in a given environment based on pre-existing parental gene expression differences.

### Homoeolog expression bias is consistent and mirrors expression differences of the diploid progenitor species

We observed a significant expression bias toward the *Brassica oleracea* (BnC) subgenome in all six resynthesized lines. The consistency of the observed bias suggests that subgenome expression dominance is likely determined based on pre-existing genetic differences between the parental genomes. Expression differences observed in the six resynthesized lines are also observed in the comparison of the diploid progenitor species. This further bolsters the claim that subgenome expression dominance is inherited from progenitors rather than an outcome of interspecific hybridization and whole genome duplication (Buggs et al. 2014). However, given that 7 of 16 individuals had significantly deviation from parental expectations of proportion of BnC biased homoeologs there appears to be some variability in the extent to which homoeologous pairs are biased toward the BnC subgenome. An excess of BnC biased pairs is also reflective of the pattern identified in natural *Brassica napus* cultivar Darmor-bzh (Chalhoub et al. 2014), but runs counter to observations of more BnA biased pairs genome-wide and in genes related to cyto-nuclear interactions in other resynthesized *Brassica napus* lines (Wu et al. 2018; Ferreira de Carvalho et al. 2019). The difference between the results presented here are likely, in part, due to *B. rapa* (BnA) being used as the maternal parent in these other two studies. Furthermore, differences in methodology may have impacted the results obtained in these other studies. For example, we control for homoeologous exchanges that alter gene dosage by excluding genomic regions that are not in a proper 2:2 ratio between both parents. Gene dosage is known to produce dosage dependent expression changes (Conant et al. 2014). Furthermore, the differences between our study and these two previous studies ((Wu et al. 2018; Ferreira de Carvalho et al. 2019) could, in part, be due to genetic differences in the accessions used to resynthesize *Brassica napus*. Our model of subgenome dominance predicts subgenome dominance is related to methylated TE density differences near genes (Bird et al. 2018). Gene and TE content are known to be highly variable within a species (Golicz et al. 2016; Anderson et al.). Due to this intraspecific variation, the progenitor genomes may differ in such a way to yield opposite subgenome dominance patterns in different resynthesized allopolyploid lines.

Gene ontology and pathway enrichment analyses revealed that BnC biased homoeologs are highly enriched with primary metabolic and organellar functions, while *Brassica rapa* (BnA) biased homoeologs were not enriched with any known biological functions. Additionally the protein-protein interaction network constructed from BnC biased genes are more highly interconnected and clustered in the network compared to BnA biased genes. These results suggest that BnC biased homoeologs are to a greater extent regulating vital cellular functions. The enrichment of organellar functions towards the BnC subgenome, which is the maternally contributed subgenome in our resynthesized lines, may be due to imprinting, but it is not definitive until similar analyses are repeated with reciprocal crosses. (Wu et al. 2018; Ferreira de Carvalho et al. 2019) (2019) analyzed 110 nuclear encoded components of plastid protein complexes and failed to find maternal progenitor homoeolog expression bias in resynthesized *Brassica napus*. Of these 110 genes, we found 41 biased toward the BnC subgenome in all six lines in generation one, 24 in generation five, and 14 in generation ten. None of the 110 genes were biased toward the BnA subgenome in all six lines in any generation. However, because our analyses are genome-wide rather than just the 110 gene subset involved in plastid complexes, we were able to identify a BnC subgenome bias towards the organelles based on the thousands of known nuclear encoded organellar genes (Savage et al. 2013) as indicated by the 263 BnC biased genes with GO cellular component annotation for the chloroplast.

### Homoeolog methylation patterns are consistent, are correlated with observed expression biases, and partially reflect differences in progenitors

We also identified consistent DNA methylation pattern differences between subgenomes. These differences appear to be a combination of inherited progenitor methylation patterns and methylation changes following interspecific hybridization and polyploidization. BnC biased homeologs showed markedly lower CG methylation at the transcription start site and lower CHG and CHH methylation in the gene body compared to BnA biased homoeologs. These DNA methylation patterns are each associated with higher gene expression (Niederhuth et al. 2016). Globally, CG, CHG, and CHH methylation of genes and LTR retrotransposons is higher in the dominant BnC subgenome than the BnA subgenome. In mimulus, a similar DNA methylation pattern was observed for the dominantly expressed subgenome (15). These methylation patterns were also present when comparing diploid progenitor epigenomes, again suggesting that this aspect of subgenome dominance is largely due to inherited features from the diploid progenitor genomes. However, it remains unclear whether these methylation pattern differences are the causal mechanism or are possibly just the result of expression differences of biased homoeologs. This may be a promising new avenue to explore to further our understanding of the underlying mechanisms of subgenome expression dominance.

While methylation differences between biased homoeolog pairs and global subgenome differences appear to be inherited from the diploid progenitor genomes, we also identified methylation patterns that appear to be a response to interspecific hybridization and/or polyploidy. CHH methylation of both BnA and BnC genes and LTR retrotransposons showed transgressive hypermethylation patterns, first matching parental methylation levels in the first generation, and then surpassing parental methylation levels in fifth and tenth generations. This increase of LTR methylation over parental and mid-parent value in later generations matches previous observations in resynthesized *B. napus* that LTR derived 21 and 24nt siRNAs are transgressively expressed in later generations (Martinez Palacios et al. 2019). Additionally for the BnC subgenome, CG methylation also appeared transgressively hypermethylated in the later generations. The functional impact, if any, of hypermethylation at CG sites on driving subgenome expression dominance remains poorly understood and should be the focus of future studies.

We found that replaying the “evolutionary tape” with resynthesized polyploids resulted in several findings relevant to further dissecting the underlying mechanisms driving subgenome expression dominance. Collectively, results from this study suggest that subgenome expression dominance is largely predetermined based on pre-existing parental gene expression differences. While the observed DNA methylation differences between subgenomes reflects expression bias, the causal role of DNA methylation patterns, particularly the pre-existing differences among the diploid progenitor genomes, in impacting subgenome expression differences remains poorly understood. Future progress in these areas will bring us closer to answering looming questions on the causes of subgenome dominance and the connections between subgenome expression and DNA methylation differences in polyploid genomes.

## Materials & Methods

### Plant growth, tissue collection, library prep

The resynthesized B. napus allopolyploid lines (CCAA) were obtained from a previous study (Xiong et al. 2011). Plants were grown at 23C during day and 20C at night with 16hr days in a growth chamber. True leaf three was collected from all plants within one hour starting at 10am (4hrs into day) and immediately flash-frozen in liquid nitrogen. Leaves were split in half for RNA and DNA isolation. Total RNA and DNA was isolated using the respective KingFisher Pure Plant kits (Thermo Fisher Scientific, MA) and quantified using Qubit 3 Fluorometer (Thermo Fisher Scientific, MA). DNA and RNA libraries were prepared using the KAPA HyperPrep and mRNA HyperPrep kit protocol, respectively (KAPA Biosystems, Roche, USA). Bisulfite conversion was performed using the EZ DNA Methylation Kit (Zymo, CA) All libraries were submitted to a genomics facility (Beijing Nuohe Zhiyuan Technology Co., Beijing, China) and sequenced with paired-end 150bp reads on an Illumina HiSeq 4000 system.

### *in silico* reference genome construction

Paired end 150bp genomic illumina reads for the doubled haploid *Brassica rapa* accession IMB-218, were aligned to the *Brassica rapa* R500 reference genome using bowtie2 v. 2.3.4.1 (Langmead and Salzberg 2012) on default settings with the flag “--very-sensitive-local”. The resulting alignment files were sorted and had read groups added with PicardTools v. 2.8.1 and SNPs were called between the R500 reference and the IMB-218 alignment using GATK v 3.5.0 Unified Genotyper, filtered to only include homozygous SNPs, and a new fasta reference was made using GATK v 3.5.0 FastaAlternativeReferenceMaker. This IMB-218 reference genome was concatenated to the TO1000 reference genome to create an *in silico* reference genome for *B. napus*.

### Homoeologous exchange analysis

Paired end 150bp genomic illumina reads were filtered with Trimmomatic v 0.33 (Bolger et al. 2014) to remove illumina TruSeq3 adapters. Trimmed reads were aligned to the *in silico B.napus* reference genome with Bowtie2 v.2.3.4.1(Langmead and Salzberg 2012) on default settings with the flag “--very-sensitive-local”. Bam files sorted with bamtools (Barnett et al. 2011) for use in downstream analyses.

MCScan toolkit (Tang et al. 2008) was used to identify syntenic, homologous gene pairs (syntelogs) between *Brassica rapa* (reference genome R500) and *Brassica oleracea* (reference genome TO1000;30). In the synthetic polyploid these can be thought of as syntenic homoeologs. Bed files based on chromosome and start/stop position information for each subgenome were generated. For all 18 samples (6 individuals x 3 generations) read depth for the A subgenome (BnA) syntenic homoeologs was determined in Bedtools (Quinlan and Hall 2010) with BedCov using the R500 syntelog bed file and for the C subgenome (BnC) using the TO1000 syntelog bed file. In R v 3.4.1, read depths for each syntenic homoeolog was normalized to reads per million for subgenome of origin and the ratio of reads mapping to a syntenic homoeolog compared to the overall read mapping for a syntenic homoeolog pair was averaged over a window of 50 genes with a step of one gene.

Homoeologous exchanged regions were identified by calculating average read depth for the BnC subgenome along a sliding window of 170 (85 up- and down stream) genes and step size of one. If 10 or more consecutive genes had a read depth within a pre-selected range it was called a homoelogous exchange. Regions 0 ≤ read depth < 0.2 were predicted to be in a 0BnC-to-4BnA ratio, 1BnC-to-3BnA was predicted for 0.2 ≤ read depth < 0.4, 2BnC-to-2BnA was predicted for 0.4 ≤ read depth < 0.6, 3BnC-to-1BnA for read depth between 0.6 ≤ read depth <0.8 and 4BnC-to-0BnA for read depth between 0.8 ≤ read depth < 1. HEs were plotted with the R package Rideogram (Hao et al.)

### RNASeq analysis

raw RNA-seq reads were filtered with Trimmomatic v 0.33 (Bolger et al. 2014)to remove illumina TruSeq3 adapters and mapped to the *in silico* reference using STAR v 2.6.0 (Dobin et al. 2013) on default settings. Transcripts were quantified in transcripts per million (TPM) from RNAseq alignments using StringTie v 1.3.5 (Pertea et al. 2015). Because the syntelogs in the progenitor genomes are in the subgenomes of the synthetic polyploids, they can be thought of as syntenic homoeologs. To avoid dosage imbalance only syntenic homoeologs determined to be at a 2:2 dosage balance were analyzed for homeolog expression bias. Additionally, to remove lowly expressed genes that might be noise, syntenic homoeologs were only kept if the total TPM of the pair was greater than 10. Syntenic homoeolog pairs with Log2 Foldchange greater than 3.5 were called BnC biased, and less than 3.5 were called BnA biased. Because lack of subgenome dominance would follow a normal distribution where deviations from 0 FC is equal in either direction, a Chi-squared goodness of fit test was carried out to test for normality. The R package Upsetr was used to identify and plot syntenic homoeologs shared by all lines for a given generation. For eeach generation, *Arabidopsis thaliana* orthologs were identified for genes showing the same subgenome bias in all six lines and the progenitors and were investigated for GO and KEGG pathway enrichment (Ashburner et al. 2000; Kanehisa and Goto 2000) in the STRING PPI network (Szklarczyk et al. 2017) using the online STRING network search application. STRING also calculated and reported average node degree, clustering coefficients, and enrichment for network interactions.

### DNA methylation analysis

The genomes of both *B. oleracea* TO1000 (EnsemblPlants 43(Parkin et al. 2014; Kersey et al. 2018) and *B. rapa* R500 were combined into a single fasta file to create an *in silico* allopolyploid genome. Whole genome bisulfite sequencing (WGBS) was mapped to the combined genome using methylpy v1.3.8 (Schultz et al. 2015) (see Supplementary Table X); using cutadapt v2.3 (Martin 2011) for adaptor trimming, Bowtie2 v2.3.5 (Langmead and Salzberg 2012) for alignment, and Picard tools v2.20.2 for marking duplicates. The chloroplast genome is unmethylated in plants and can be used as in internal control for calculating the non-conversion rate of bisulfite treatment (percentage of unmethylated sites that fail to be converted to uracil)(Lister et al. 2008). In mapping the parental TO1000 and R500 genomes, a small fraction of reads (∼1.3% TO1000 and ∼6.1% IMB218) mapped to the wrong genome, but seems to have had a limited impact on overall results.It is likely that there is a small percentage of mismapping in the resynthesized allopolyploids as well. As an additional control for this, we randomly down-sampled TO1000 to an equal number of read pairs as IMB218. These were combined with the IMB218 reads and mapped to the combined genome to mimic an *in silico* allopolyploid. By including this *in silico* allopolyploid alongside the individually mapped parents, we should be able to observe the influence of mismapping. DNA methylation levels in this combined dataset were either approximately half-way between the two parents for the whole genome or at a sub-genome level, approximately equal to that of the respective parent. This indicates that mismapping has little effect on genome-wide analyses, although it may still be a factor at specific regions.

Genome-wide levels of DNA methylation and DNA methylation metaplots were analyzed as previously described (Niederhuth et al. 2016) using python v3.7.3. Pybedtools v (Dale et al. 2011) and Bedtools v2.25.0 (Quinlan and Hall 2010). Briefly, for genome-wide DNA methylation levels, the weighted methylation level(Schultz et al. 2012), which accounts for sequencing coverage, was calculated for each sequence context (CG, CHG, and CHH). This was done for the combined genome, and each individual subgenome. For gene metaplots, cytosines from 2 kbps upstream, 2 kbs downstream and within the gene body were extracted. Cytosines within gene bodies were restricted to those found in coding sequences, as the presence of TEs in introns and problems of proper UTR annotation can obscure start/stop sites and introduce misleadingly high levels of DNA methylation (Niederhuth et al. 2016). Each of these three regions were then divided into 20 windows and the weighted methylation level for each window calculated and average for all genes. For LTR metaplots, the same analysis was performed, except the all cytosines within the LTR body were included. Metaplots were created for all genes in the combined genome, all genes within each subgenome, syntenic genes in each subgenome, and non-syntenic genes in each subgenome. All plots and statistics were done in R v3.6.0 (Anon), plots made using ggplot2 (Wickham 2009). All code and original analyzed data and plots are available on Github (https://github.com/niederhuth/Replaying-the-evolutionary-tape-to-investigate-subgenome-dominance).

## Data availability

Raw data from this project is available on the NCBI Sequence Read Archive (SRA) Project XXX

## Supplemental Information

**Figure S1.**
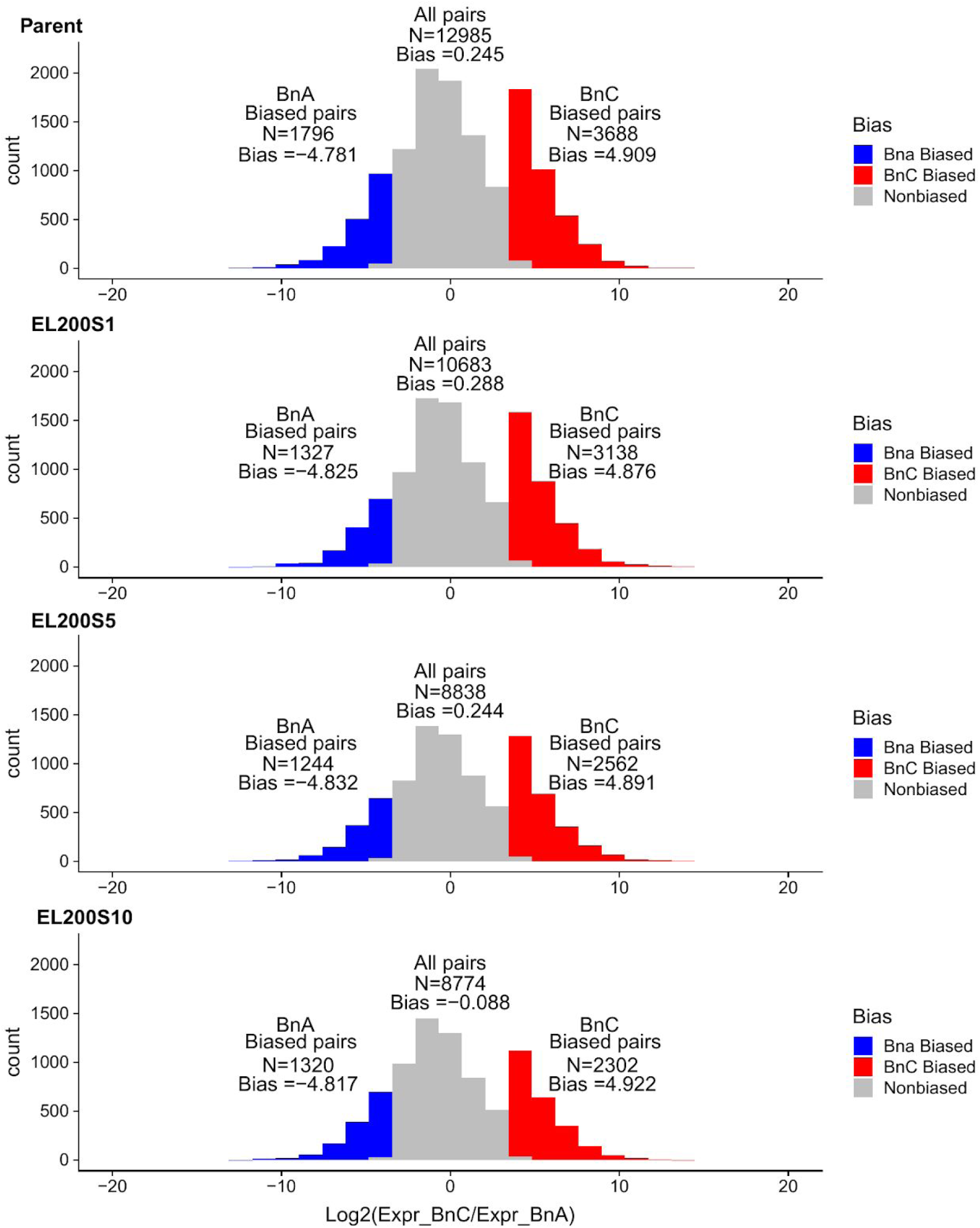
**Homoeolog Expression Bias:** Distribution of homoeolog bias in the parent and three generations of line EL 200, EL 300, EL 400, EL 600, and EL 1100, red regions indicate BnC biased homeologos with log2 expression foldchange greater than 3.5 and blue regions indicate BnA biased homoeologs with log2 expression foldchange less than −3.5.

**Figure S2.**
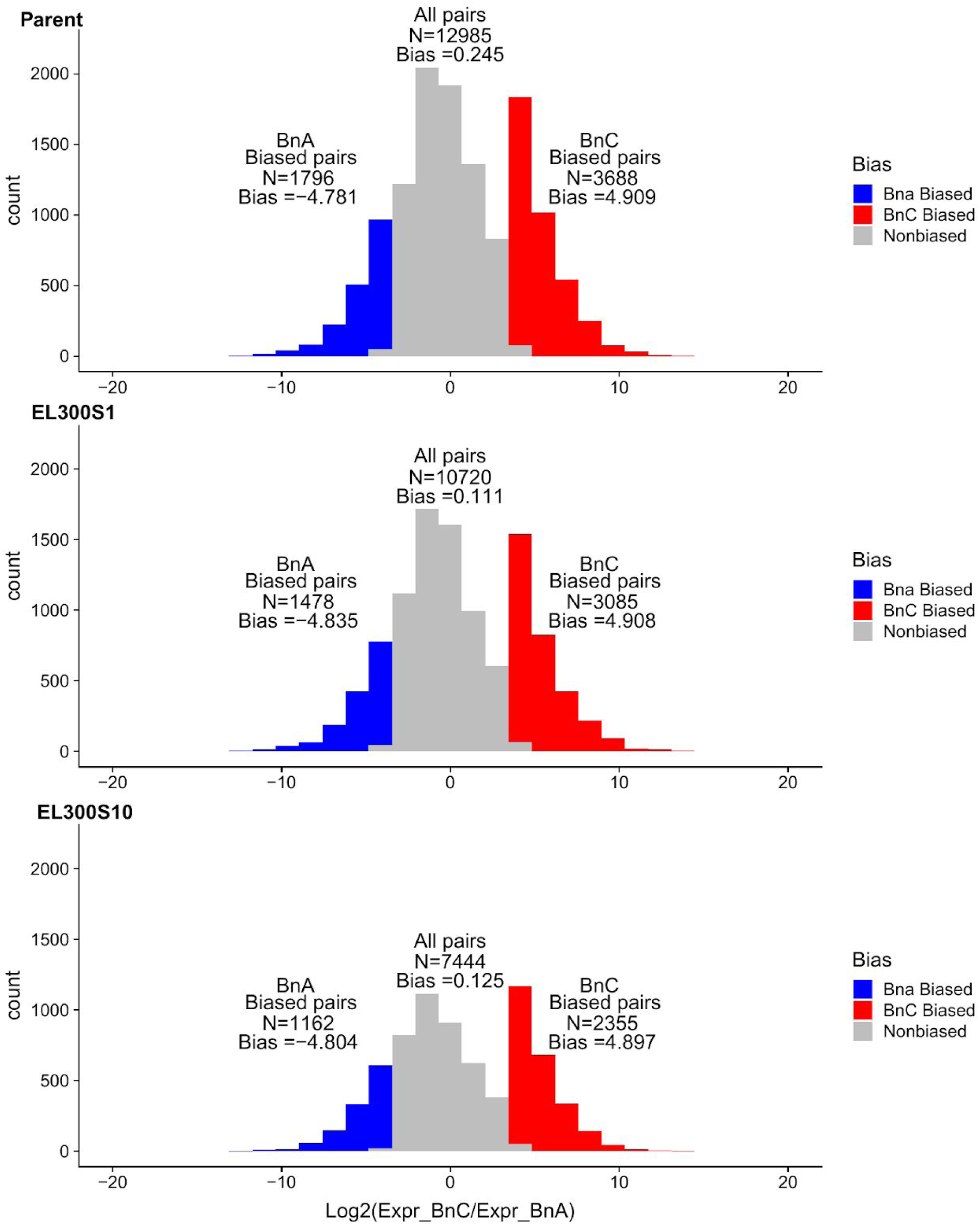
**Homoeolog Expression Bias:** Distribution of homoeolog bias in the parent and three generations of line EL 200, EL 300, EL 400, EL 600, and EL 1100, red regions indicate BnC biased homeologos with log2 expression foldchange greater than 3.5 and blue regions indicate BnA biased homoeologs with log2 expression foldchange less than −3.5.

**Figure S3.**
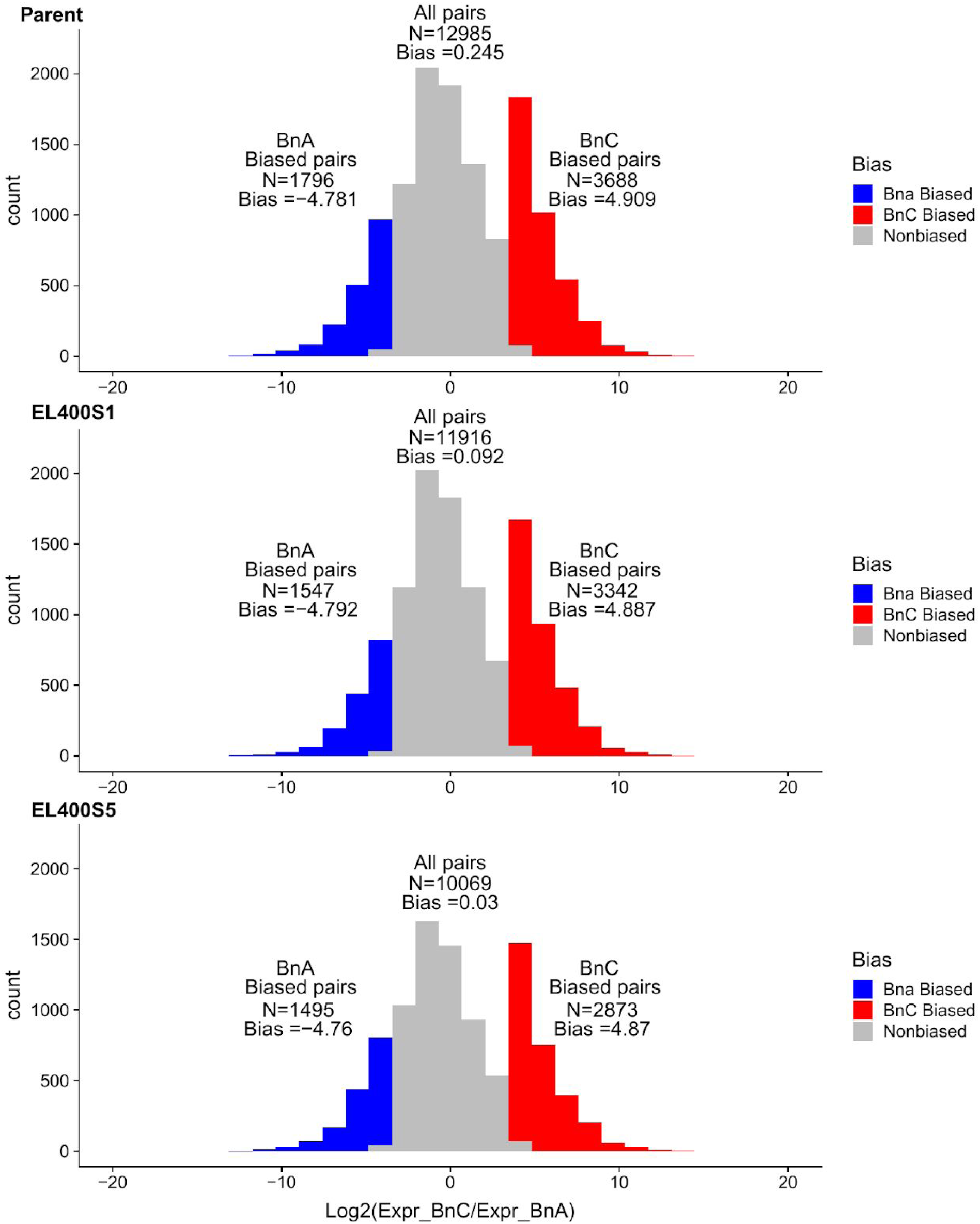
**Homoeolog Expression Bias:** Distribution of homoeolog bias in the parent and three generations of line EL 200, EL 300, EL 400, EL 600, and EL 1100, red regions indicate BnC biased homeologos with log2 expression foldchange greater than 3.5 and blue regions indicate BnA biased homoeologs with log2 expression foldchange less than −3.5.

**Figure S4.**
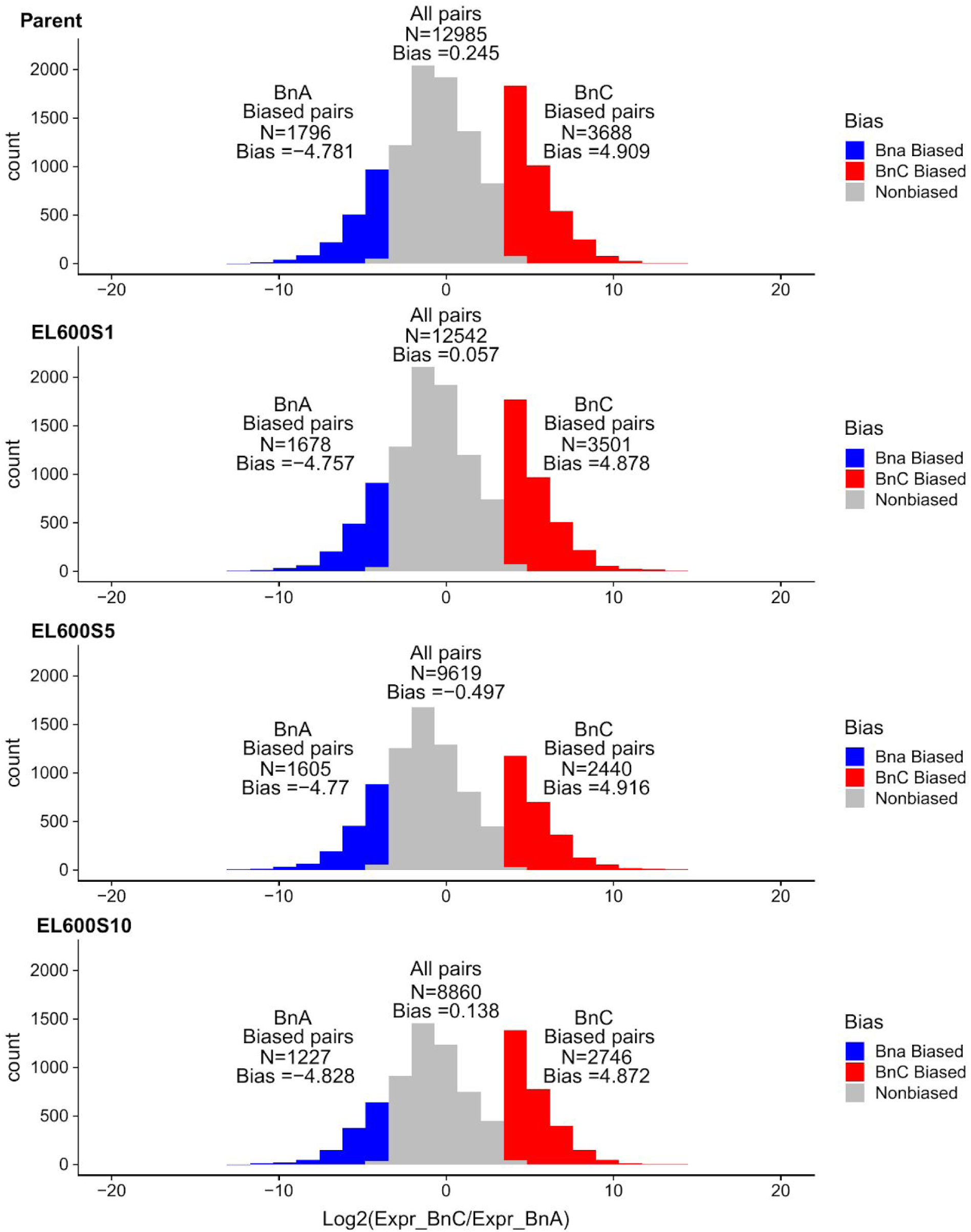
**Homoeolog Expression Bias:** Distribution of homoeolog bias in the parent and three generations of line EL 200, EL 300, EL 400, EL 600, and EL 1100, red regions indicate BnC biased homeologos with log2 expression foldchange greater than 3.5 and blue regions indicate BnA biased homoeologs with log2 expression foldchange less than −3.5.

**Figure S5.**
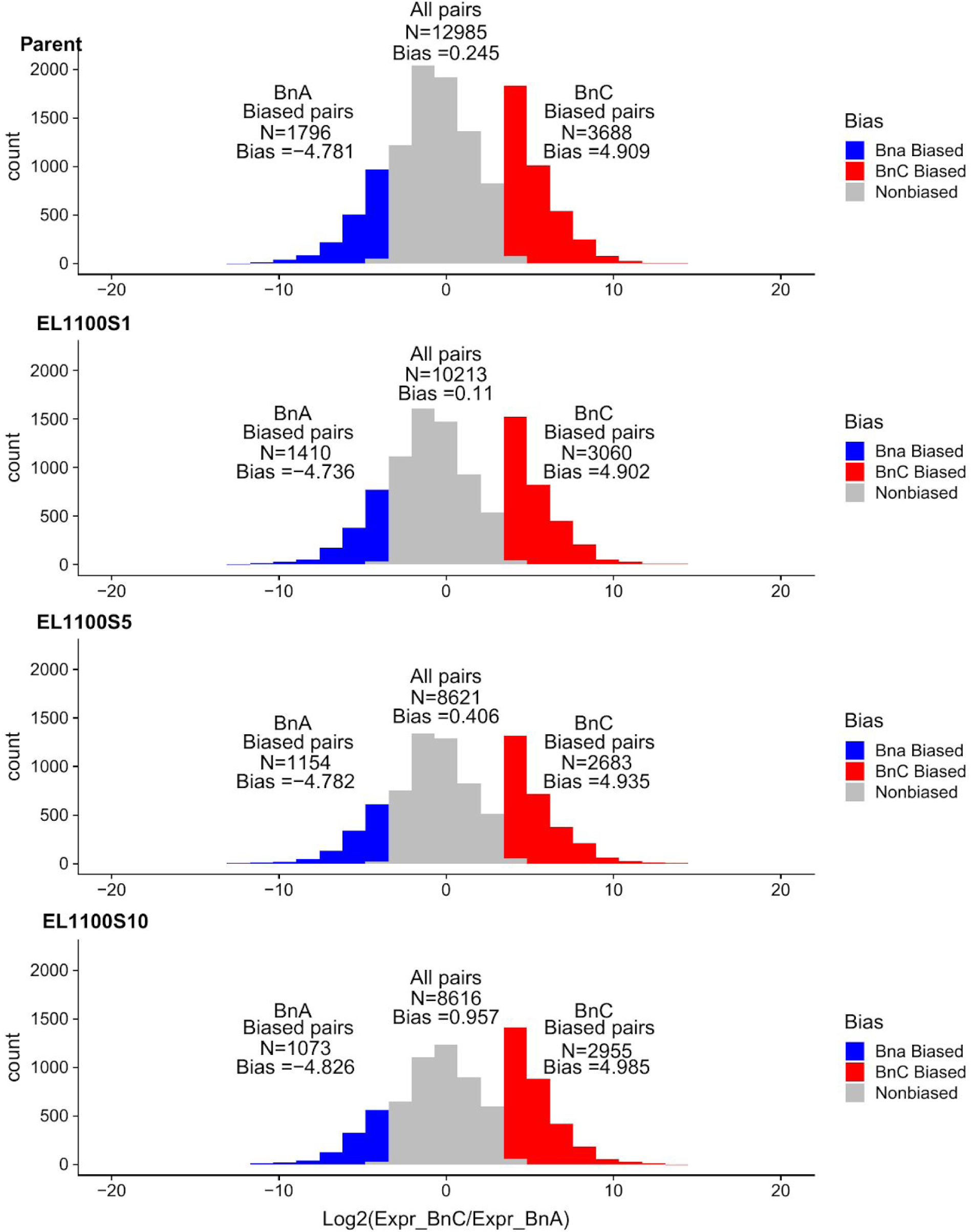
**Homoeolog Expression Bias:** Distribution of homoeolog bias in the parent and three generations of line EL 200, EL 300, EL 400, EL 600, and EL 1100, red regions indicate BnC biased homeologos with log2 expression foldchange greater than 3.5 and blue regions indicate BnA biased homoeologs with log2 expression foldchange less than −3.5.

**Figure S6.**
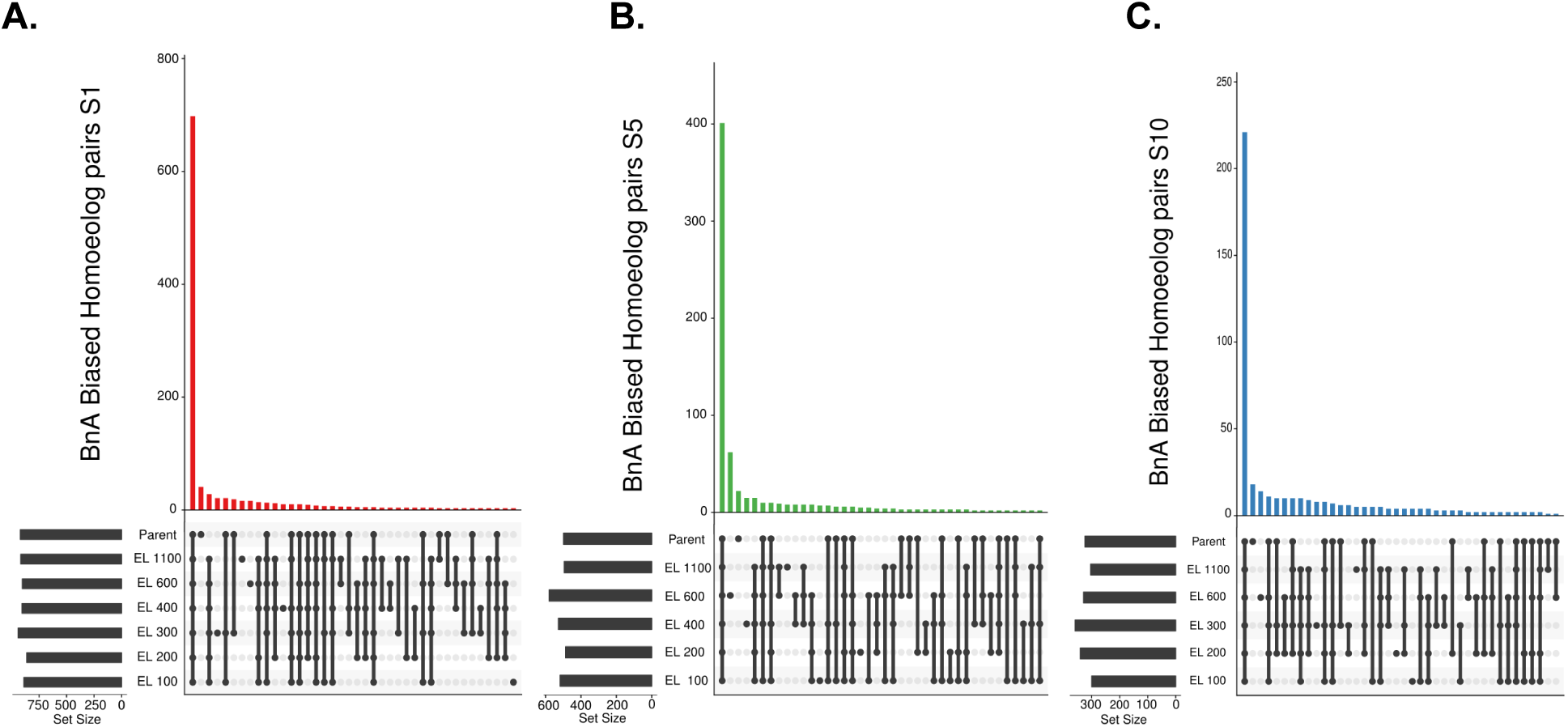
Common shared biased Homoeolog pairs: Upset plot of how biased homoeologs pairs for BnA biased (**a-c)** are shared among all six lines for the three sampled generations. These analysis was restricted only to homoeolog pairs in 2:2 balance in all 6 lines

**Figure S7.**
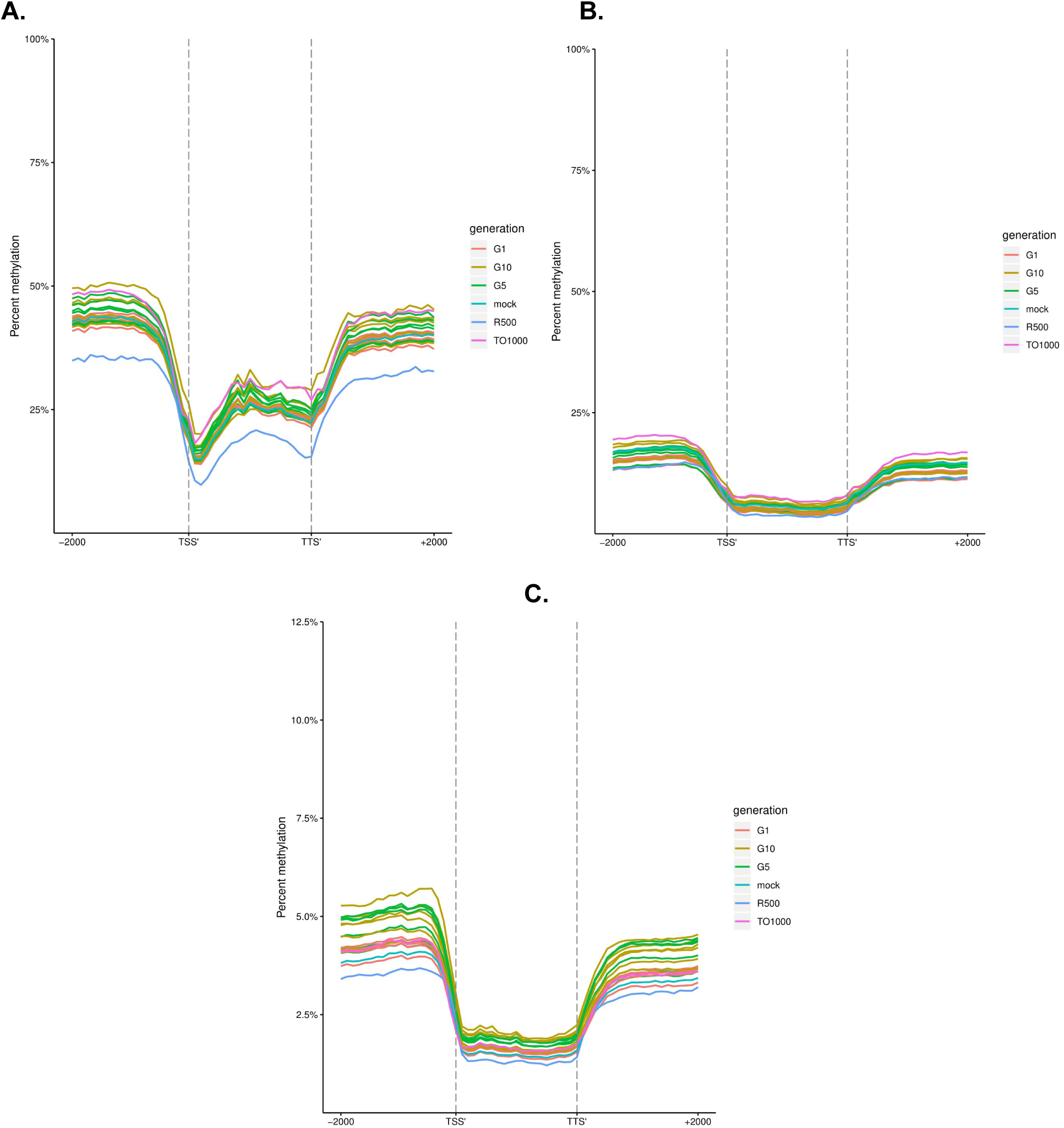
Metaplots of CG (**a**), CHG (**b**), and CHH (**c**) mean weighted methylation of all annotated gene models 2kb upstream of the transcription start site, gene body, the transcription termination site, and 2kb downstream of the transcription termination site for both parents, an *in silico* “mock” polyploid and the three generations of the resynthesized polyploids for all six lines

**Figure S8.**
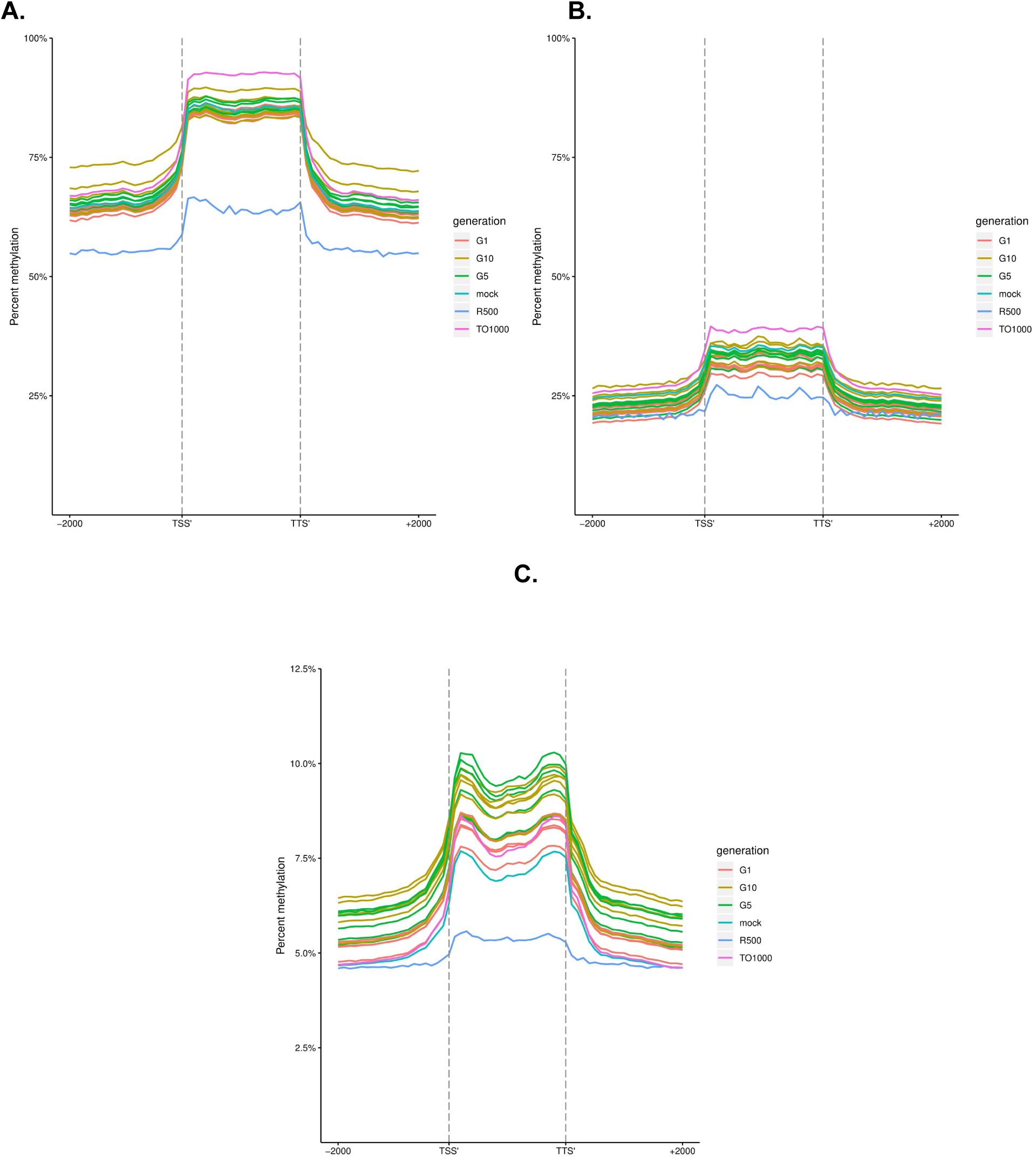
Metaplots of CG (**a**), CHG (**b**), and CHH (**c**) mean weighted methylation of all annotated Long tandem repeat transposable elements 2kb upstream of the transcription start site, gene body, the transcription termination site, and 2kb downstream of the transcription termination site for both parents, an *in silico* “mock” polyploid and the three generations of the resynthesized polyploids for all six lines

**Figure S9.**
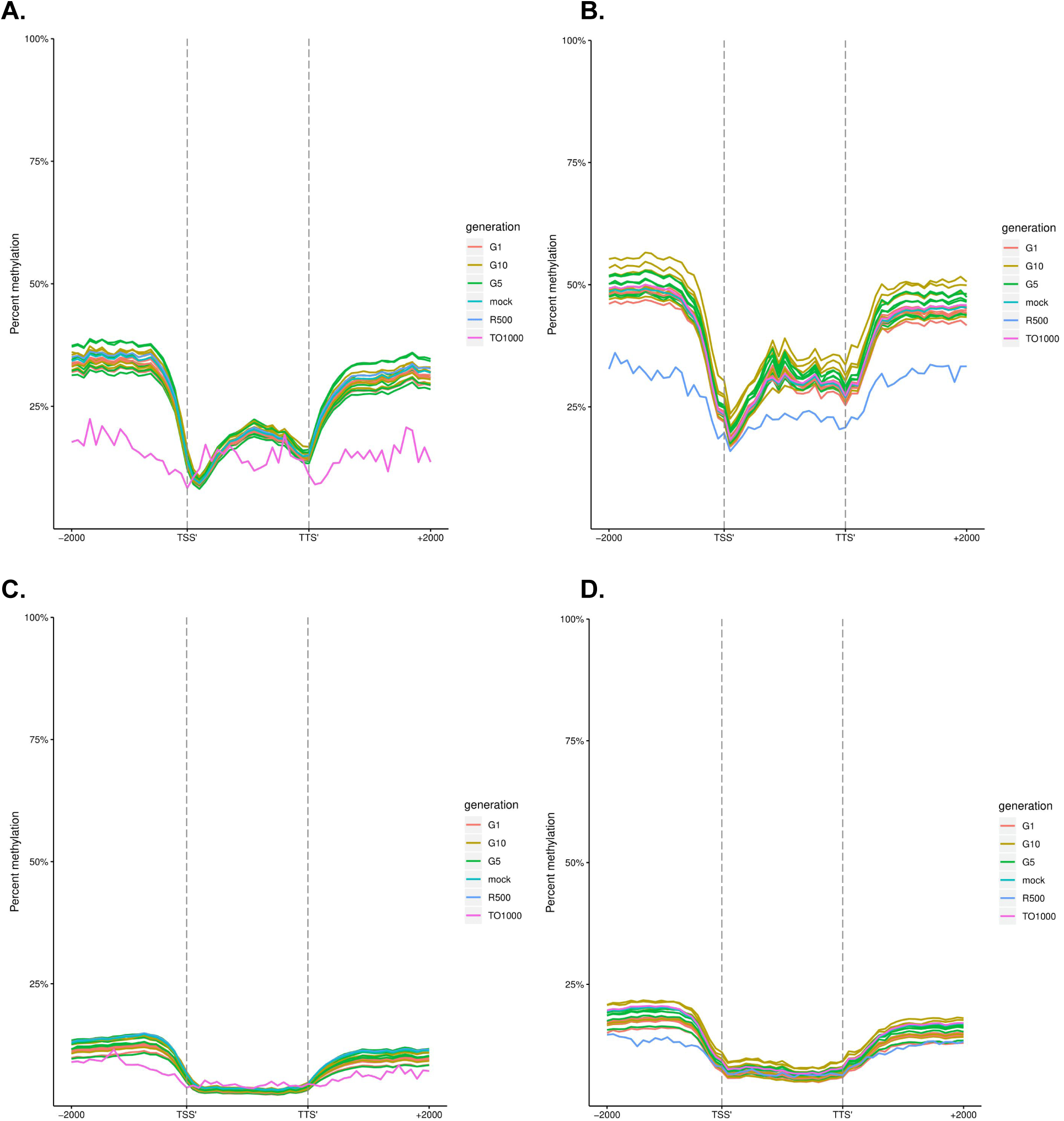
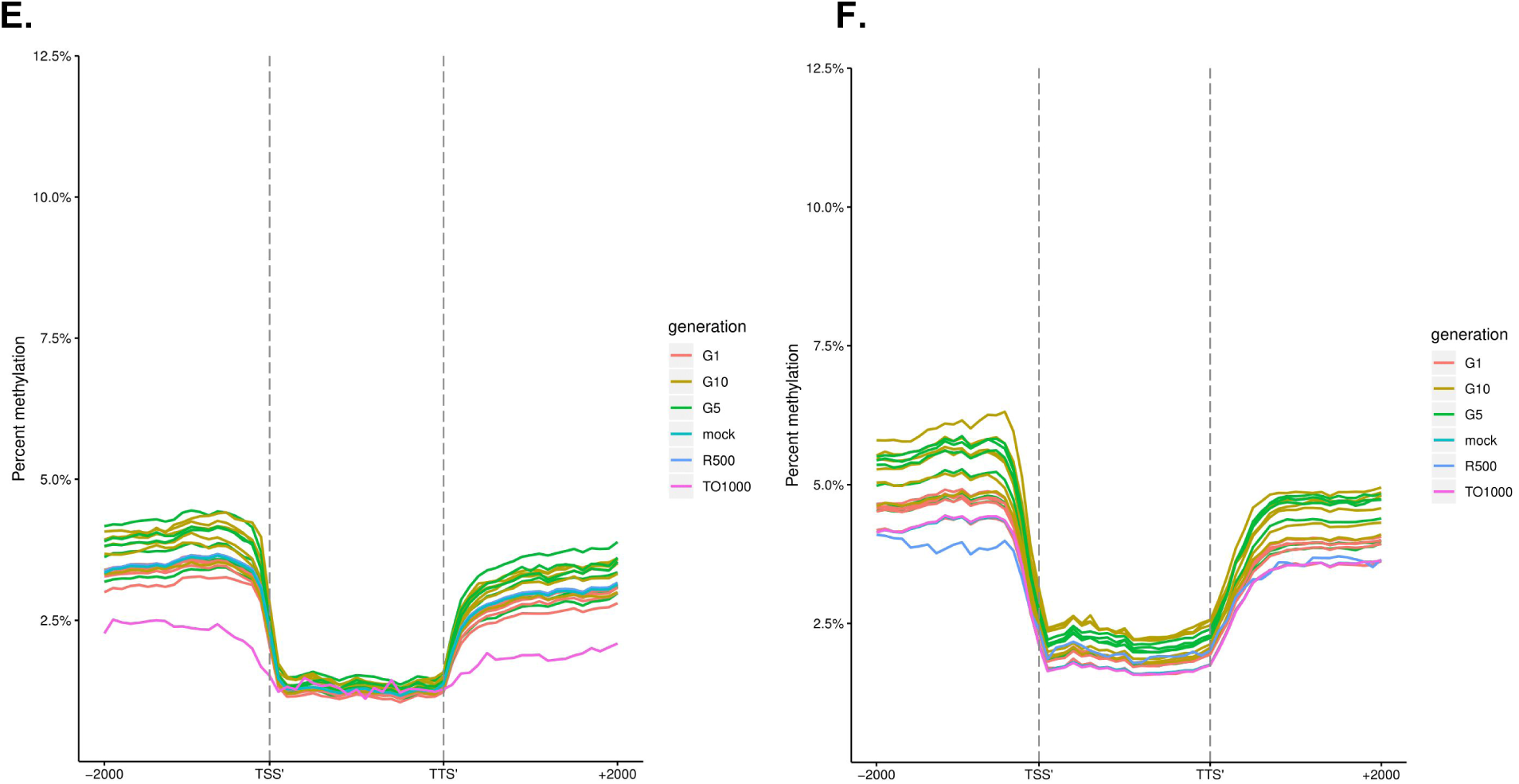
a-c Metaplots of CG (**a,b)**, CHG (**c,d)**, and CHH (**e,f**) mean weighted methylation of all annotated A (**a,c,e)** and C (**b,d,f**) subgenome gene models 2kb upstream of the transcription start site, gene body, the transcription termination site, and 2kb downstream of the transcription termination site for both parents, an *in silico* “mock” polyploid and the three generations of the resynthesized polyploids for all six lines

**Figure S10.**
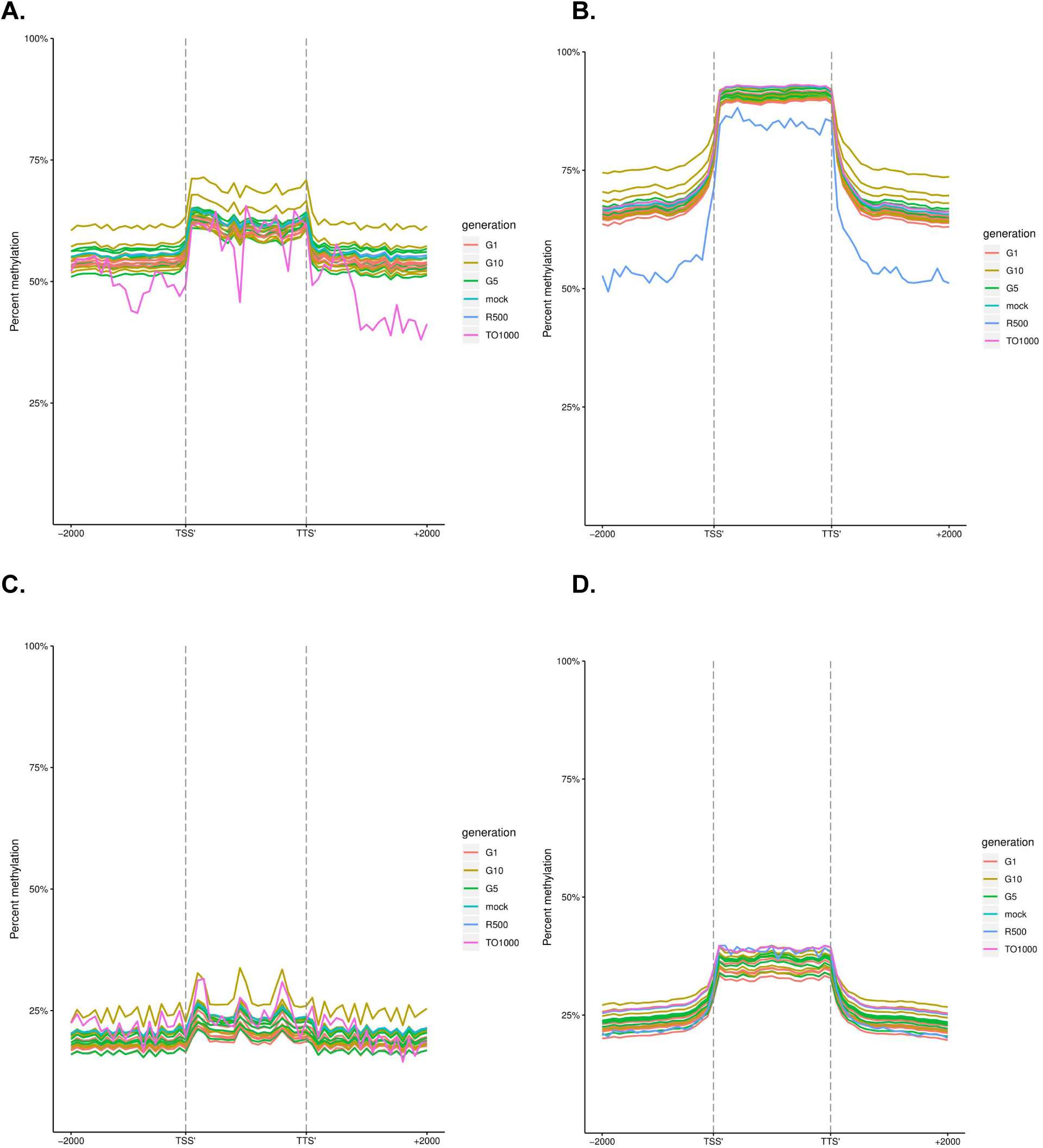
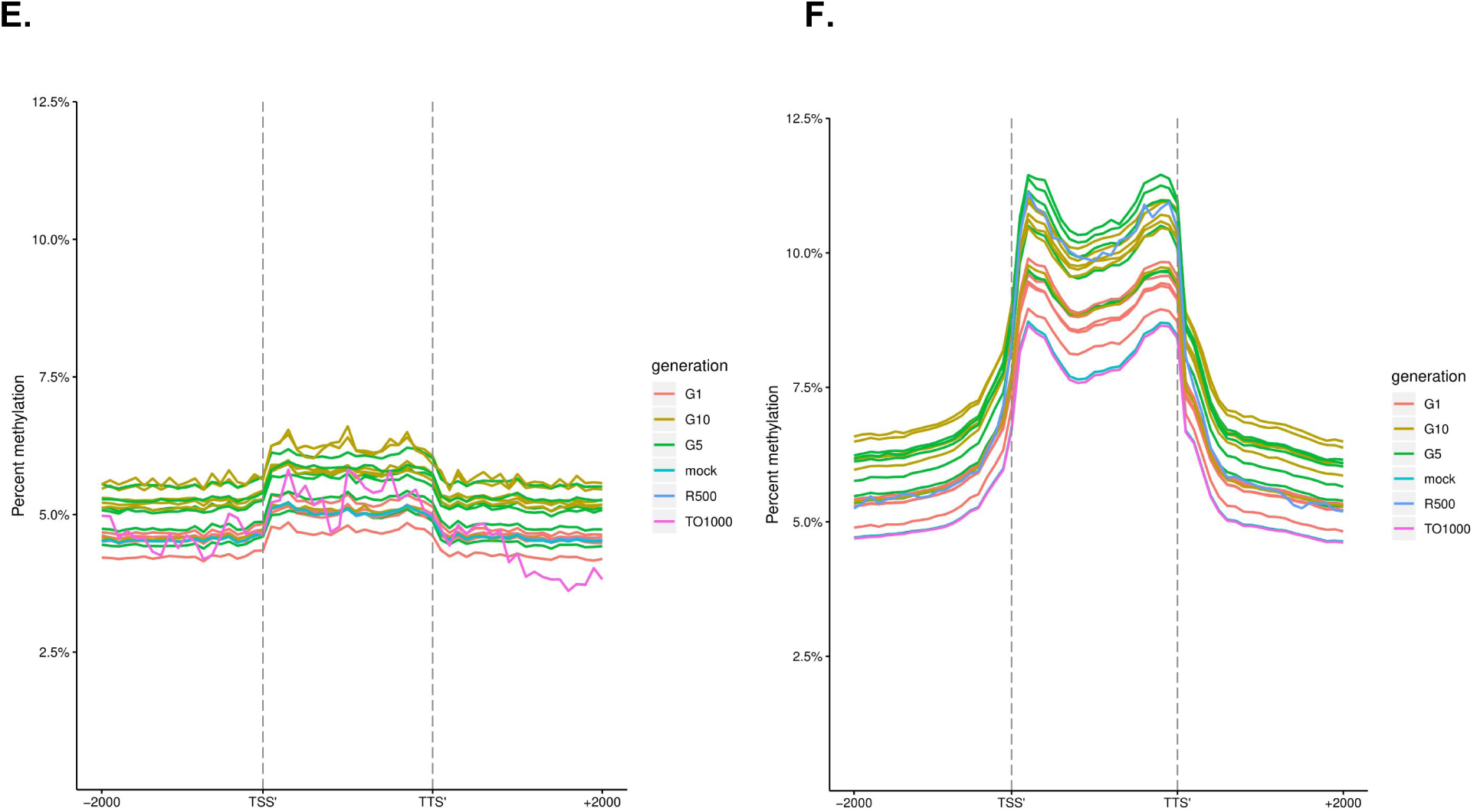
Metaplots of CG (**a,b)**, CHG (**c,d)**, and CHH (**e,f**) mean weighted methylation of all annotated A (**a,c,e)** and C (**b,d,f**) subgenome long tandem repeat transposable elements 2kb upstream of the transcription start site, gene body, the transcription termination site, and 2kb downstream of the transcription termination site for both parents, an *in silico* “mock” polyploid and the three generations of the resynthesized polyploids for all six lines

**Figure S11.**
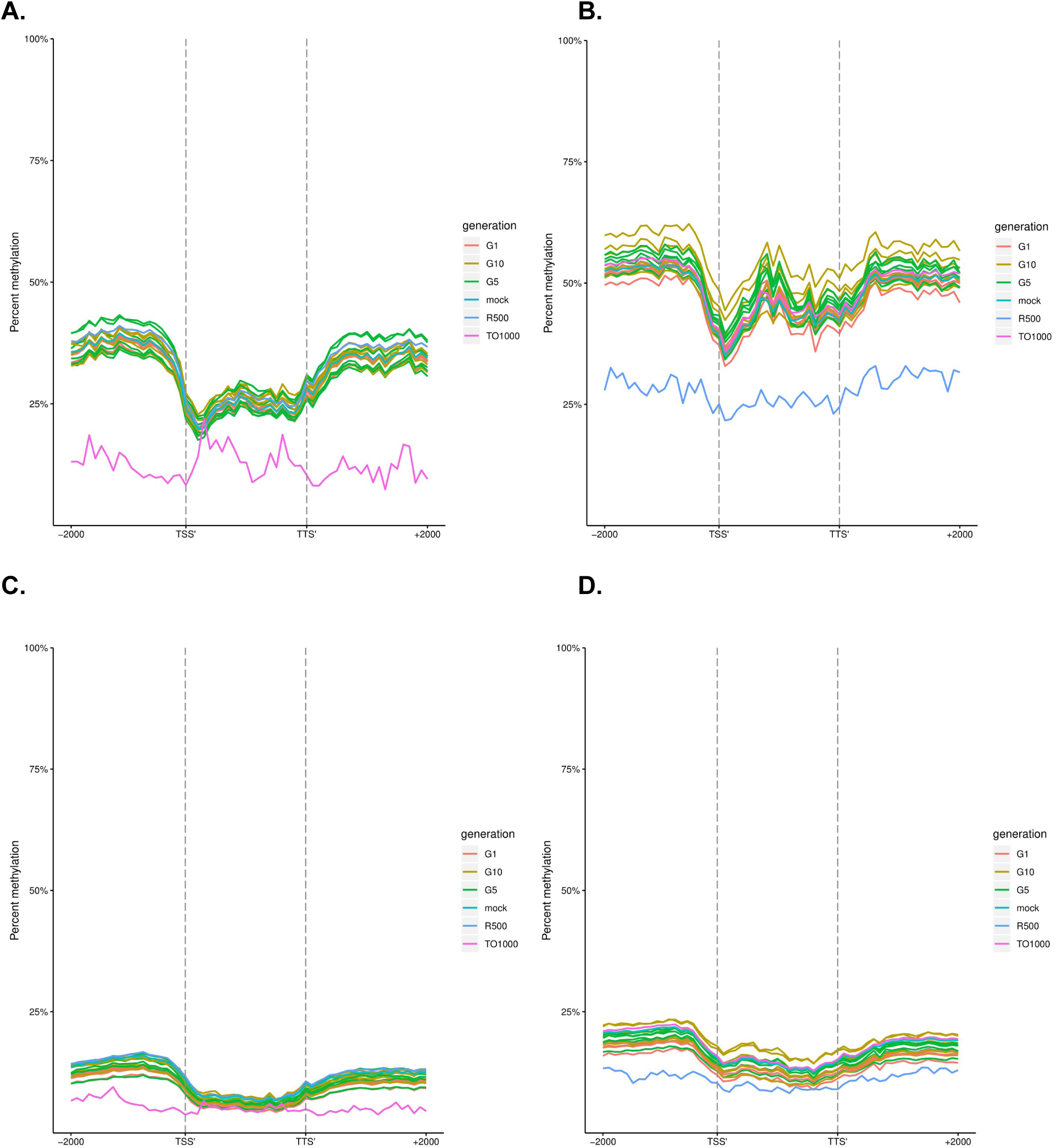
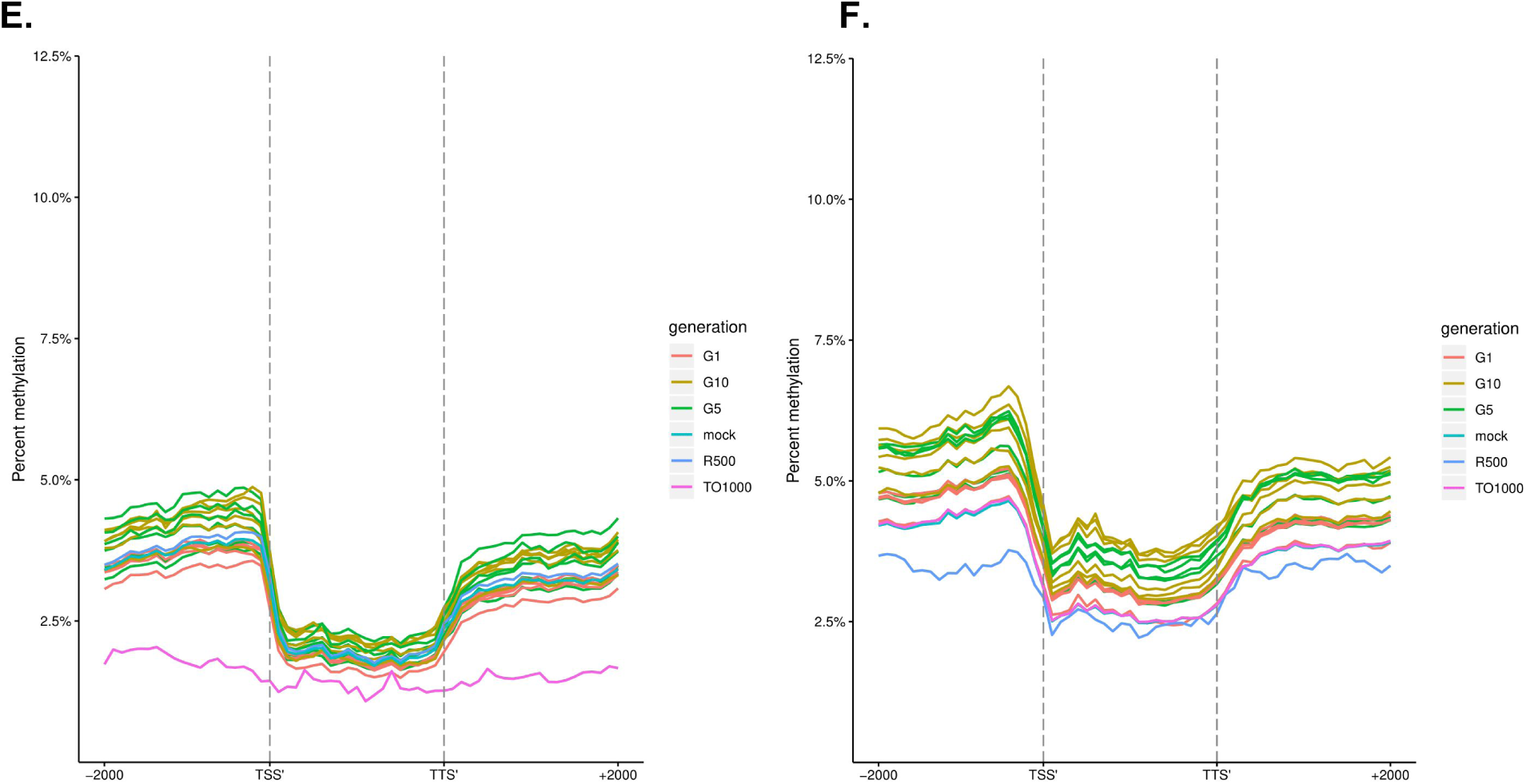
Metaplots of CG (**a,b)**, CHG (**c,d)**, and CHH (**e,f**) mean weighted methylation of non-syntenic A (**a,c,e)** and C (**b,d,f**) subgenome gene models 2kb upstream of the transcription start site, gene body, the transcription termination site, and 2kb downstream of the transcription termination site for both parents, an *in silico* “mock” polyploid and the three generations of the resynthesized polyploids for all six lines

**Figure S12.**
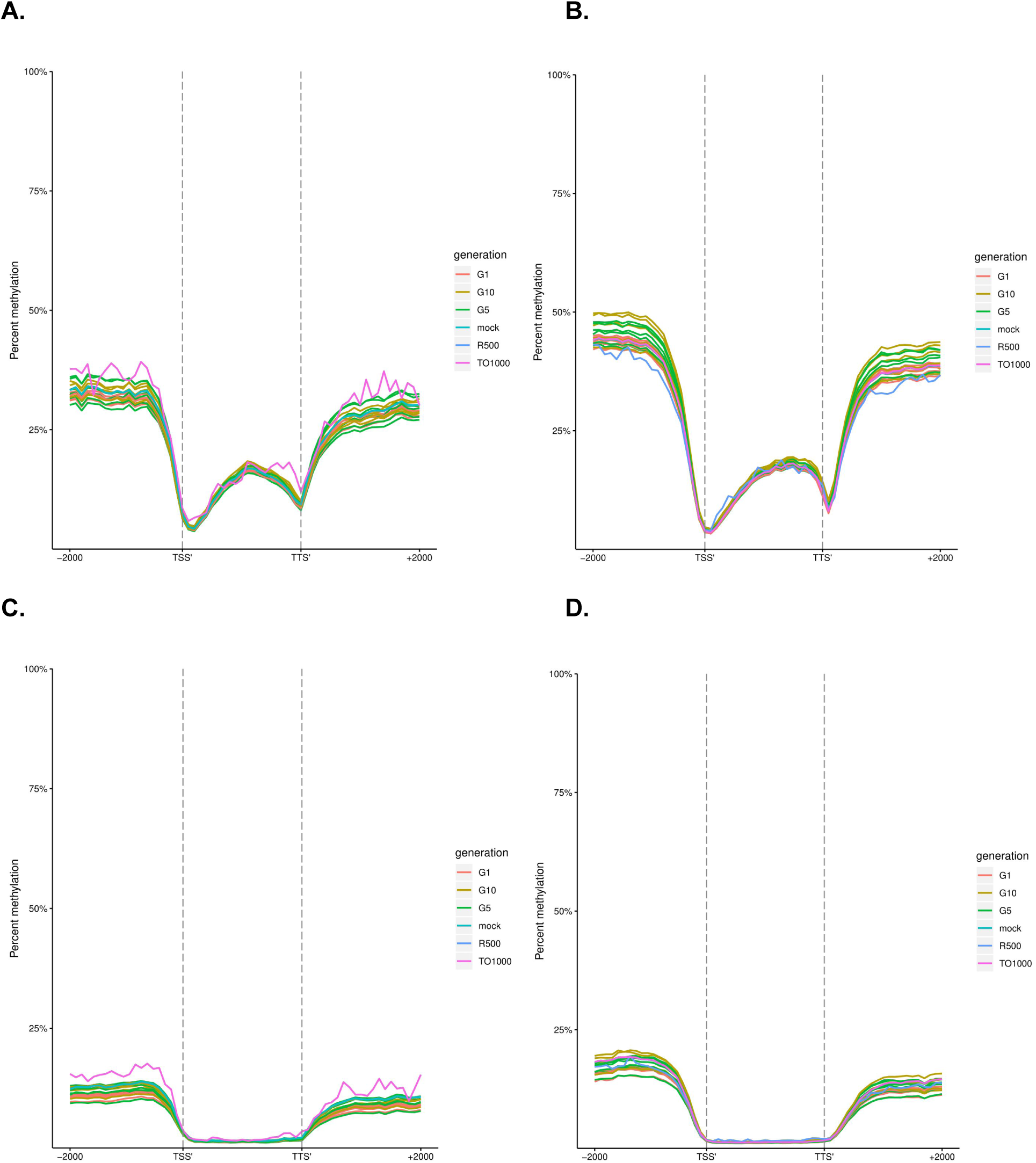
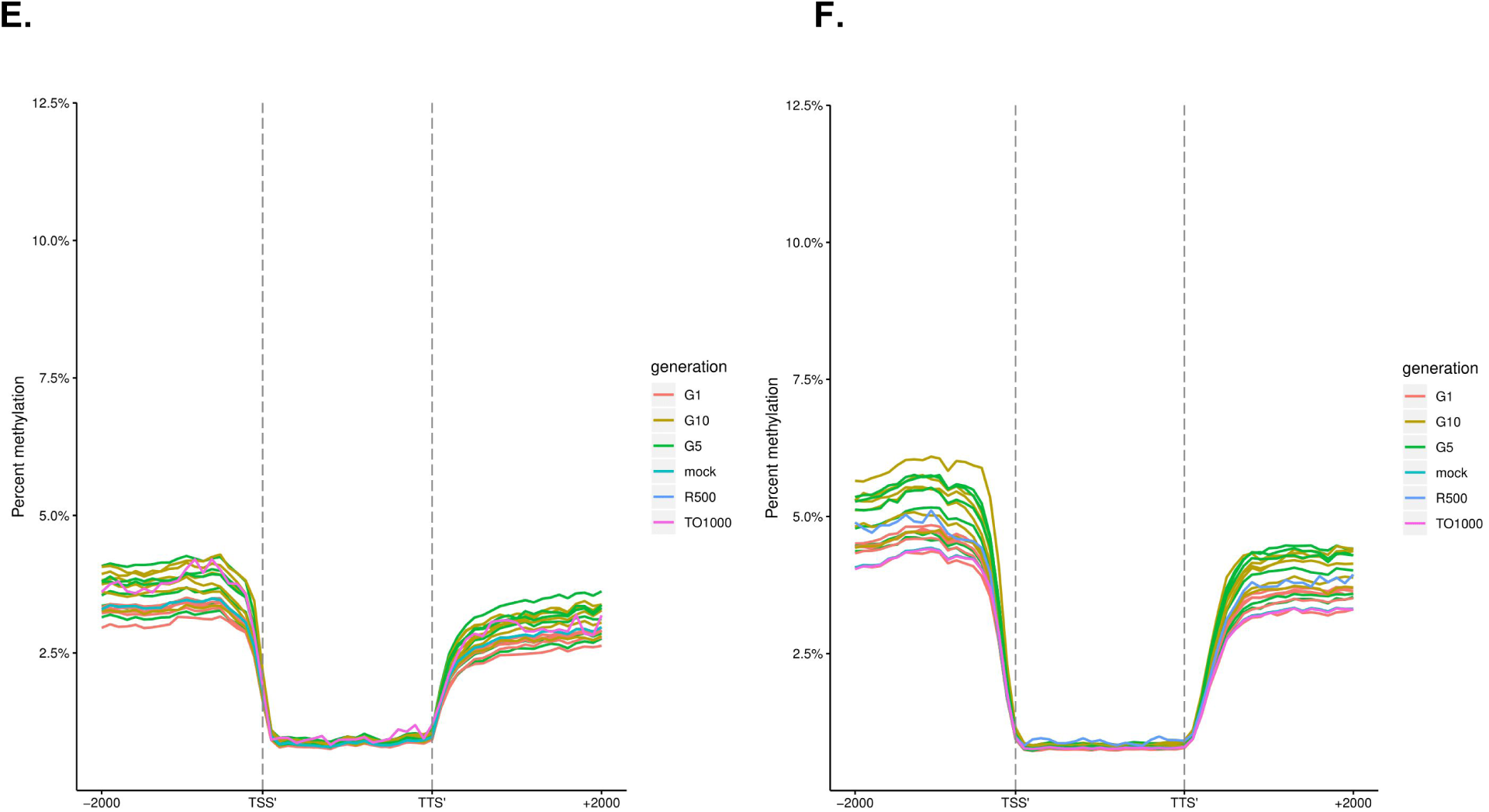
Metaplots of CG (**a,b)**, CHG (**c,d)**, and CHH (**e,f**) mean weighted methylation of syntenic A (**a,c,e)** and C (**b,d,f**) subgenome gene models 2kb upstream of the transcription start site, gene body, the transcription termination site, and 2kb downstream of the transcription termination site for both parents, an *in silico* “mock” polyploid and the three generations of the resynthesized polyploids for all six lines

**Table S1:**
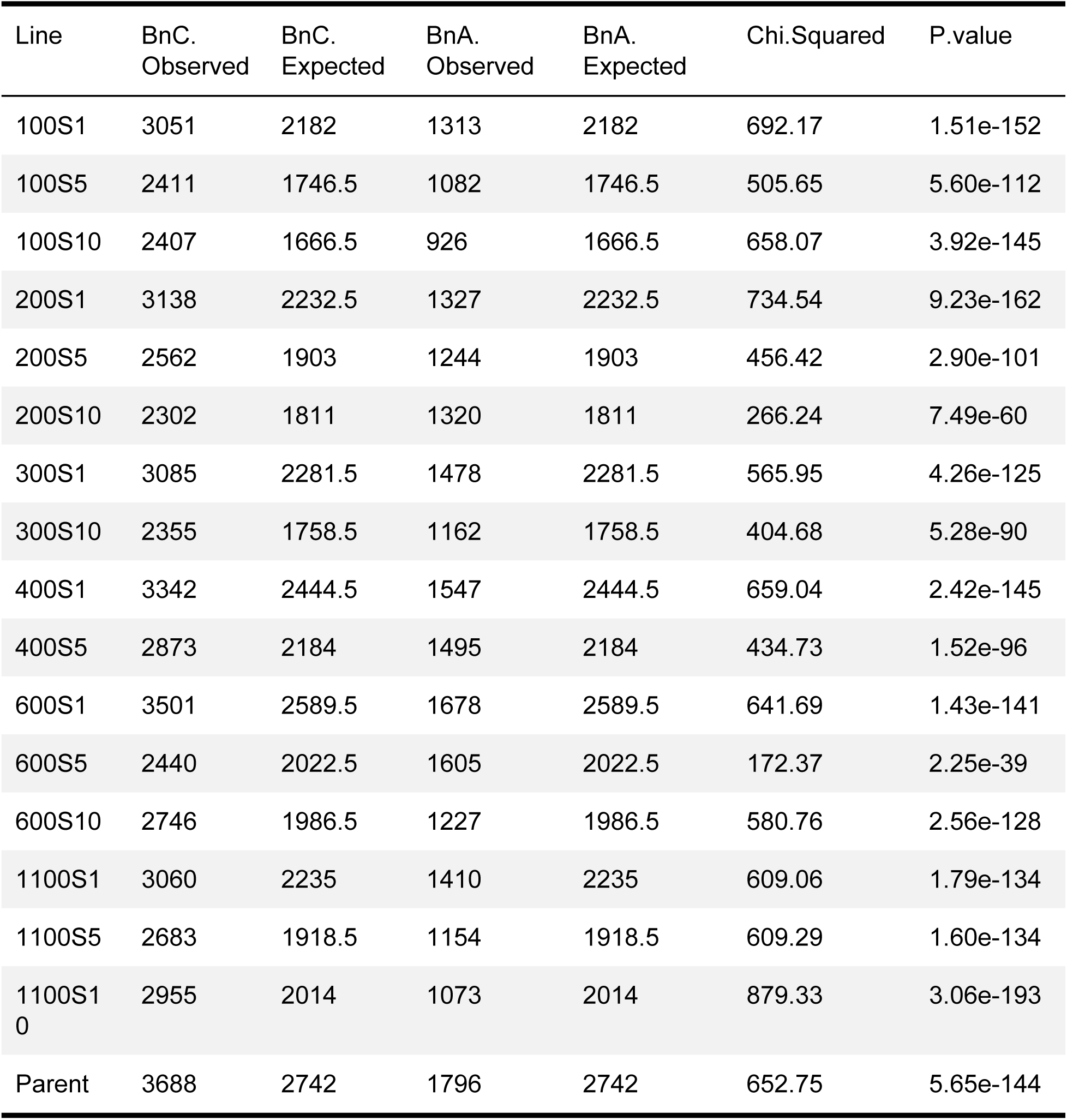
Homeolog Expression Bias Chi Squared table

**Table S2.** Homeolog Expression Bias vs Parent Chi Squared table

| Line | BnC<br>Observed | BnC<br>Expected | BnA<br>Observed | BnA<br>Expected | Chi.Squared | P.value |
| --- | --- | --- | --- | --- | --- | --- |
| 100S1 | 3051 | 2934.798 | 1313 | 1429.202 | 14.049 | 0.0002 |
| 100S5 | 2411 | 2349.049 | 1082 | 1143.951 | 4.9888 | 0.03 |
| 100S10 | 2407 | 2241.448 | 926 | 1091.552 | 37.336 | 9.94e-10 |
| 200S1 | 3138 | 3002.721 | 1327 | 1462.279 | 18.61 | 1.60e-05 |
| 200S5 | 2562 | 2559.542 | 1244 | 1246.458 | 0.0072085 | 0.9323 |
| 200S10 | 2302 | 2435.802 | 1320 | 1186.198 | 22.443 | 2.17e-06 |
| 300S1 | 3085 | 3068.626 | 1478 | 1494.374 | 0.26679 | 0.6055 |
| 300S10 | 2355 | 2365.189 | 1162 | 1151.811 | 0.13402 | 0.7143 |
| 400S1 | 3342 | 3287.861 | 1547 | 1601.139 | 2.722 | 0.09897 |
| 400S5 | 2873 | 2937.488 | 1495 | 1430.512 | 4.3229 | 0.0376 |
| 600S1 | 3501 | 3482.887 | 1678 | 1696.113 | 0.28763 | 0.5917 |
| 600S5 | 2440 | 2720.27 | 1605 | 1324.73 | 88.172 | < 2.2e-16 |
| 600S10 | 2746 | 2671.85 | 1227 | 1301.15 | 6.2836 | 0.01219 |
| 1100S1 | 3060 | 3006.083 | 1410 | 1463.917 | 2.9528 | 0.08573 |
| 1100S5 | 2683 | 2580.389 | 1154 | 1256.611 | 12.459 | 0.00042 |
| 1100S10 | 2955 | 2708.837 | 1073 | 1319.163 | 68.305 | < 2.2e-16 |

